# Aridity drives global convergence of desert microbiomes and biogeochemical activities

**DOI:** 10.1101/2025.11.29.691269

**Authors:** Pok Man Leung, Sean K. Bay, Wei Wen Wong, Thanavit Jirapanjawat, Stephen D. J. Archer, Julian Beaman, Ameur Cherif, Steven L. Chown, Don A. Cowan, Cecilia Demergasso, Asunción de los Ríos, Jocelyne DiRuggiero, Bo Elberling, Beat Frey, Osnat Gillor, David W. Graham, Puja Gupta, Ian D. Hogg, Heli Juottonen, Minna-Maarit Kytöviita, Thulani Makhalanyane, Laura K. Meredith, Thanh Nguyen-Dinh, Anders Priemé, Jean-Baptiste Ramond, Steven D. Siciliano, Geok Yuan Annie Tan, Kimberley A. Warren-Rhodes, Nimrod Wieler, Perran L. M. Cook, Manuel Delgado-Baquerizo, Chris Greening

**Affiliations:** Department of Microbiology, Monash Biomedicine Discovery Institute, Clayton VIC 3800, Australia; School of Biological Sciences, Monash University, Clayton VIC 3800, Australia; Securing Antarctica’s Environmental Future, Monash University, Clayton VIC 3800, Australia; Water Studies, School of Chemistry, Monash University, Clayton VIC 3800, Australia; AgResearch Group, Bioeconomy Science Institute, Palmerston North 4442, New Zealand; Higher Institute of Biotechnology of Sidi Thabet (ISBST), University of Manouba, Ariana, Tunisia; Centre for Microbial Ecology and Genomics, Department of Biochemistry, Genetics and Microbiology, University of Pretoria, Pretoria 0002, South Africa; Department of Biochemistry, Genetics and Microbiology, University of Pretoria, Pretoria 0002, South Africa; Biotechnology Center, Universidad Católica del Norte, Antofagasta, Chile; Department of Biogeochemistry and Microbial Ecology, National Museum of Natural Sciences, Madrid, Spain; Department of Biology, Department of Earth and Planetary Sciences, Johns Hopkins University, Baltimore, Maryland, USA; Center for Permafrost, Department of Geosciences and Natural Resource Management, University of Copenhagen; Swiss Federal Research Institute WSL, Birmensdorf, Switzerland; Zuckerberg Institute for Water Research, Blaustein Institutes for Desert Research, Ben Gurion University of the Negev, Sede Boker, Israel; School of Engineering, Newcastle University, Newcastle Upon Tyne, United Kingdom; Fermentation and Microbial Biotechnology Division, CSIR-Indian Institute of Integrative Medicine, Jammu, Jammu & Kashmir, India; Department of Biotechnology, School of Biosciences, RIMT University, Punjab, India; School of Science, University of Waikato, Hamilton, New Zealand; Polar Knowledge Canada, Canadian High Arctic Research Station, Cambridge Bay, Nunavut, Canada; Department of Biological and Environmental Science, University of Jyväskylä, Finland; School of Natural Resources and the Environment, University of Arizona, Tucson, AZ, USA; Center for Volatile Interactions, Department of Biology, University of Copenhagen, Denmark; Extreme Ecosystem Microbiomics & Ecogenomics Laboratory, Faculty of Biological Sciences, Pontificia Universidad Católica de Chile, Santiago, Chile; Department of Soil Science, University of Saskatchewan, Saskatoon, Canada; Institute of Biological Sciences, University of Malaya, Kuala Lumpur, Malaysia; Carl Sagan Center, SETI Institute and NASA Ames Research Center, Mountain View, CA, USA; Research Department, Israel Antiquities Authority, POB 586, Jerusalem 91004, Israel; Laboratorio de Biodiversidad y Funcionamiento Ecosistémico, Instituto de Recursos Naturales y Agrobiología de Sevilla (IRNAS), Consejo Superior de Investigaciones Científicas (CSIC). Av. Reina Mercedes 10, E-41012 Sevilla, Spain

**Author notes:** **Corresponding authors:** Pok Man Leung, Chris Greening.

## Abstract

Deserts cover a third of the world’s surface, supporting unique biomes and ecosystem services. Yet, we lack a comprehensive assessment of what defines and drives the microbial communities that dominate life in these regions. Here, we conducted a standardized field survey in contrasting cold, hot, and polar deserts across the seven continents, and observed geographically distant deserts share similar structure, function, and activities. Desert communities are dominated by genomically streamlined Actinobacteriota and Chloroflexota, and compared with non-desert soils, are significantly enriched with stress tolerance genes, mobile genetic elements, and antiviral strategies, revealing previously unknown ecological and evolutionary dynamics. Metabolically, these communities exhibit reduced capacity for carbohydrate and protein degradation, and instead are enriched for chemosynthetic carbon fixation, continuous energy harvesting using atmospheric trace gases and sunlight, and energy reserve biosynthesis. All sampled soils mediated respiration, trace gas oxidation, and carbon fixation, with detectable activity even in hyper-arid Atacama and Antarctic soils at the margins of life. Driver analyses identified aridity as the primary overriding driver of the microbial communities and biogeochemical activities. Collectively, these findings suggest that aridity selects for metabolically self-sufficient taxa capable of continuously meeting energy and carbon needs independently of vegetation-derived inputs, while enduring physicochemical stressors and potentially elevated viral pressure. These new insights are integral to forecast the future of soils amid increasing desertification.

**Significance statement:** Desert soils occupy a vast and expanding portion of Earth, yet what defines and governs their dominant microbial life remains incompletely defined. By assessing the composition, capabilities, and activities of microbial communities across deserts on all seven continents, we identify unifying signatures of life under extreme water limitation. We show microbial communities are highly self-sufficient, capable of acquiring energy and carbon even where plant inputs are minimal. This planetary-scale understanding of the desert microbiome has important ramifications for forecasting potential shifts of microbial communities and the services they provide as desertification intensifies.

## Introduction

Deserts cover one third of Earth’s land surfaces (1, 2). All deserts are defined by scarce and variable availability of water, compounded by stressors such as temperature extremes, ultraviolet irradiation, high salinity, and, in polar regions, extremely seasonal light variation and prolonged freezing. Collectively, these factors constrain plant cover and exclude most animals, with desert soils typically having low levels of photosynthetic primary production and in turn minimal organic carbon to sustain organoheterotrophs. Yet, abundant microbes are present in desert soils (3, 4). As aridity increases, overall microbial diversity decreases and primary producers such as Cyanobacteria decline towards undetectable levels, whereas the relative abundance of monoderm bacteria from Actinobacteriota and Chloroflexota increases (5–10). Even in the driest core of the Atacama Desert, enzymatic activity assays and metabolite profiling suggest many cells remain viable (7). What remains unclear is how desert microbes persist and sustain activity under such stressful conditions.

Desert microbes must adopt multiple strategies to endure water and energy limitations. Key physiological adaptations to desiccation include accumulating compatible solutes (e.g. trehalose, betaine, glutamate), using cations for intracellular osmoregulation (11, 12), secreting hygroscopic exopolysaccharides to retain water (13, 14), and membrane remodelling (15, 16). Stressed microbes can synthesize pigments and antioxidants, and upregulate DNA and protein repair machineries, to overcome desiccation- and UV-induced oxidative damage (17–19). The energy sources sustaining desert microbes, amid low photosynthetic inputs and high investment in stress responses, have remained obscure. It was assumed that desert microbes primarily sustain themselves by relying on organic carbon stores accumulated during transient hydration events (4). However, it was recently discovered that some bacteria can continually harvest energy and fix carbon using atmospheric trace gases such as hydrogen (H_2_), a process known as aerotrophy (8, 20–26). Some desert microbes also mediate rhodopsin-based light harvesting, sulfur oxidation, or nitrification to obtain energy (8, 21, 27, 28). Despite the progress in elucidating microbial adaptation to physical extremes, global-scale investigation of (dis)similarities and drivers of the functional microbiomes across contrasting cold, hot and polar deserts is lacking. Moreover, most insights into desert adaptations are extrapolated from non-desert model organisms and desert ecosystems are systematically underrepresented in global metagenome sampling efforts (29–31). This knowledge is critical to anticipate the future of drylands under climate change and land degradation.

Here, to address this gap, we conducted a standardized field survey of the composition, function, and activities of microbial communities in 25 hot, cold, and polar desert locations, from 15 countries and seven continents. Our study combines physicochemical profiling, gene-centric and genome-resolved metagenomics, biogeochemical and isotopic assays, and driver analyses. We identify five hallmark strategies enriched in desert microbes, namely genome streamlining and mobilization, osmolyte production, continuous energy harvesting via trace gases and sunlight, reserve compound biosynthesis, and antiviral defence, and demonstrate that aridity is the overriding driver selecting for these strategies. Such insights are of paramount importance to the prediction of community and ecosystem level response to future climate change and the consequent rapid expansion of deserts (32).

## Results and Discussion

### Geographically distant deserts select for compositionally similar communities enriched with genomically streamlined Actinobacteriota and Chloroflexota

We sampled three exposed topsoils within ten hot and five cold deserts (herein nonpolar deserts; defined by high aridity, aridity index < 0.2) and ten polar deserts (defined by low availability of non-frozen water; mean annual temperature < 0°C), mapped to IUCN desert ecoregions (2) (**Fig. 1A**). The sampled soils spanned a wide range of physicochemical characteristics (**Fig. S1**, **Table S1**). In line with previous reports (5, 8, 21), desert soils harbour numerous microbial cells (average 7.5 × 10^8^ cells per gram dry soil), though cell counts range by four orders of magnitude (from 2.8 × 10^5^ cells g_dw_^-1^ in the Atacama Desert to 2.6 × 10^9^ cells g_dw_^-1^ in Gobi Mongolia; **Table S1**).

**Figure 1.**
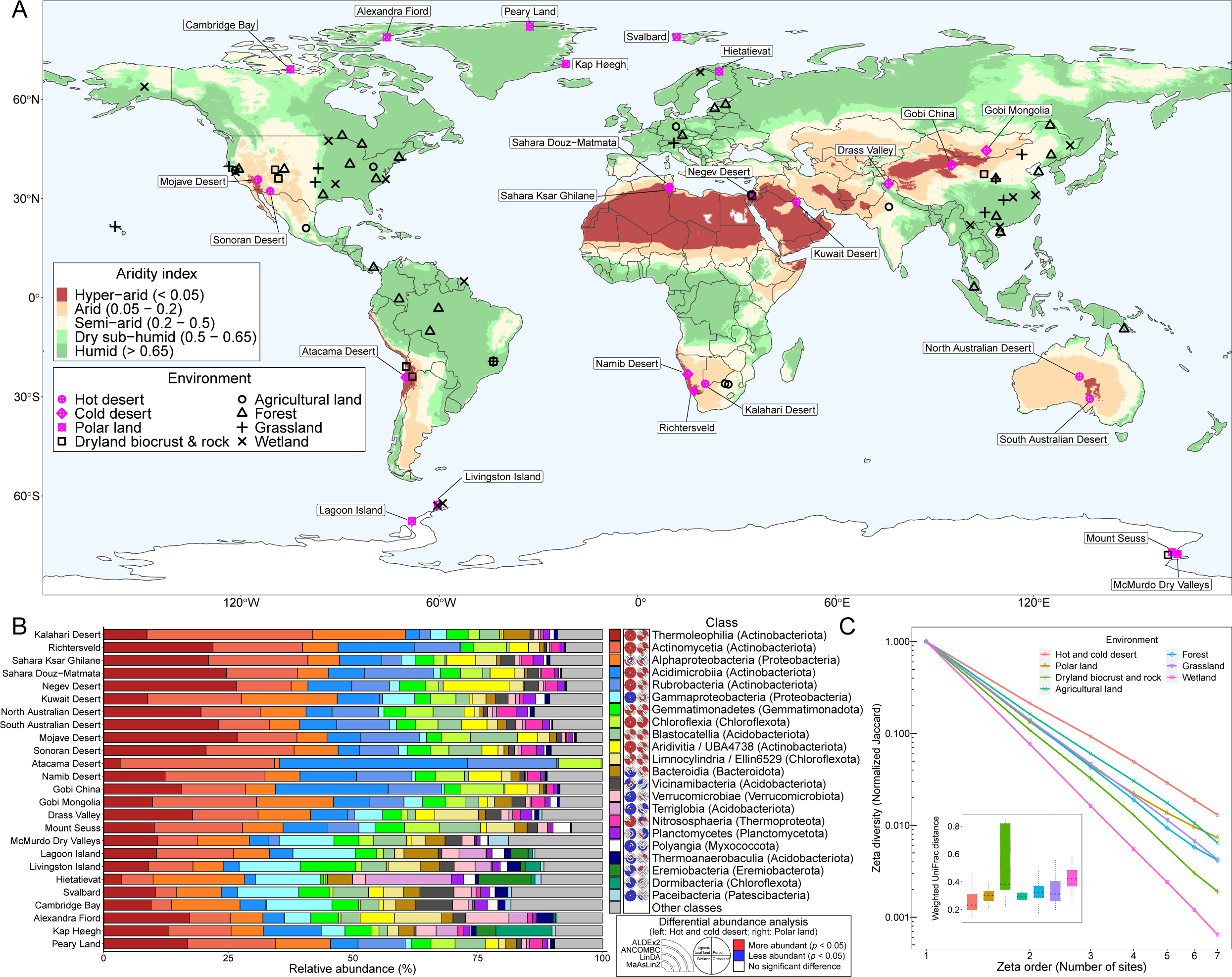
Microbial communities across twenty-five desert sampling locations. (**A**) Map displaying the global distribution of hot, cold, and polar drylands. Color shading represents the aridity index (AI), a ratio of mean annual precipitation to potential evapotranspiration. Locations of 25 sampled sites (magenta) and 67 reference soil environments (agricultural land, dryland biocrusts and rocks, forest, grassland, and wetland) (black) for comparative analysis are shown. (**B**) Average class-level composition of microbial communities from the 25 sites based on metagenomic reads of the universal single-copy marker gene *rplB*. Bacterial and archaeal taxonomy is based on the Genome Taxonomy Database (GTDB). Classes that do not attain more than 2.5% abundance in any site were grouped to “Other classes”. The donut charts show differentially abundant microbial classes in nonpolar deserts (left) and polar lands (right) compared to four non-desert soil ecosystems determined by ALDEx2, ANCOMBC, LinDA, and MaAsLin2, respectively. Significance values were adjusted by Benjamini-Hochberg procedure. (**C**) Zeta decline showing the pattern of shared richness (Jaccard normalization at genus level) when increasing number of sites from the same ecosystem category is considered. Distance decay of zeta diversity with increasing geographic distance between sites is reported in **Fig. S2H-I**. The enclosed boxplot shows pairwise community dissimilarity (beta diversity) between sites from the same ecosystem category using weighted UniFrac.

We systematically compared deeply sequenced metagenomes of all 75 sampled desert soils (**Table S2**) with those of 67 other locations (agricultural land, forest, grassland, wetland, desert biocrusts / rocks) with compatible sampling design, sequencing scheme, and geographic range (**Table S3**). The microbial community composition between these soils was compared using two independent methods (**Table S4-5**). On average, deserts comprise 94% bacteria, 1.3% archaea, and 1.1% fungi with monoderm Actinobacteriota and Chloroflexota dominating all sites and Cyanobacteria comprising just 0.1% of the community, except in the biocrust and rock samples. In contrast, non-desert sites contained higher proportions of diderm bacteria, fungi, and other eukaryotes (**Fig. 1B, Fig. S2A**). As elaborated in **Supplementary Note 1**, both the multisite incidence-based metric zeta diversity (33, 34) and pairwise abundance-based metric beta diversity show communities are much more compositionally similar within deserts across wide geographic separation compared to any other ecosystems and exhibit weaker distance decay relationships (**Fig. 1C, Fig. S2G-I**). Building on geographically restricted earlier studies (35), these findings suggest that distant deserts select for common dominant taxa.

Genome-resolved metagenomics was applied to determine traits of desert microorganisms. Reconstruction of metagenomes yielded a dereplicated set of 911 high-quality and medium-quality metagenome-assembled genomes (MAGs), covering 21 microbial phyla and each of the 30 most abundant orders in deserts (**Fig. 5A**). For comparative analysis, we also constructed 2226 new MAGs from previously sequenced public metagenomes (**Table S6**). Mean completeness-normalized genome sizes of MAGs from nonpolar deserts (3.7 Mb) and polar lands (3.8 Mb) are significantly smaller than from biocrust and rock (4.9 Mb), agricultural land (5.1 Mb), forest (5.6 Mb), grassland (4.8 Mb), and wetland (4.1 Mb) (all *p* < 0.05, Welch’s t-test) (**Fig. 2A**). This suggests that, similar to observations in the oligotrophic open ocean, desert microorganisms may be selected for with streamlined genomes and hence lower resource needs (36–39).

**Figure 2.**
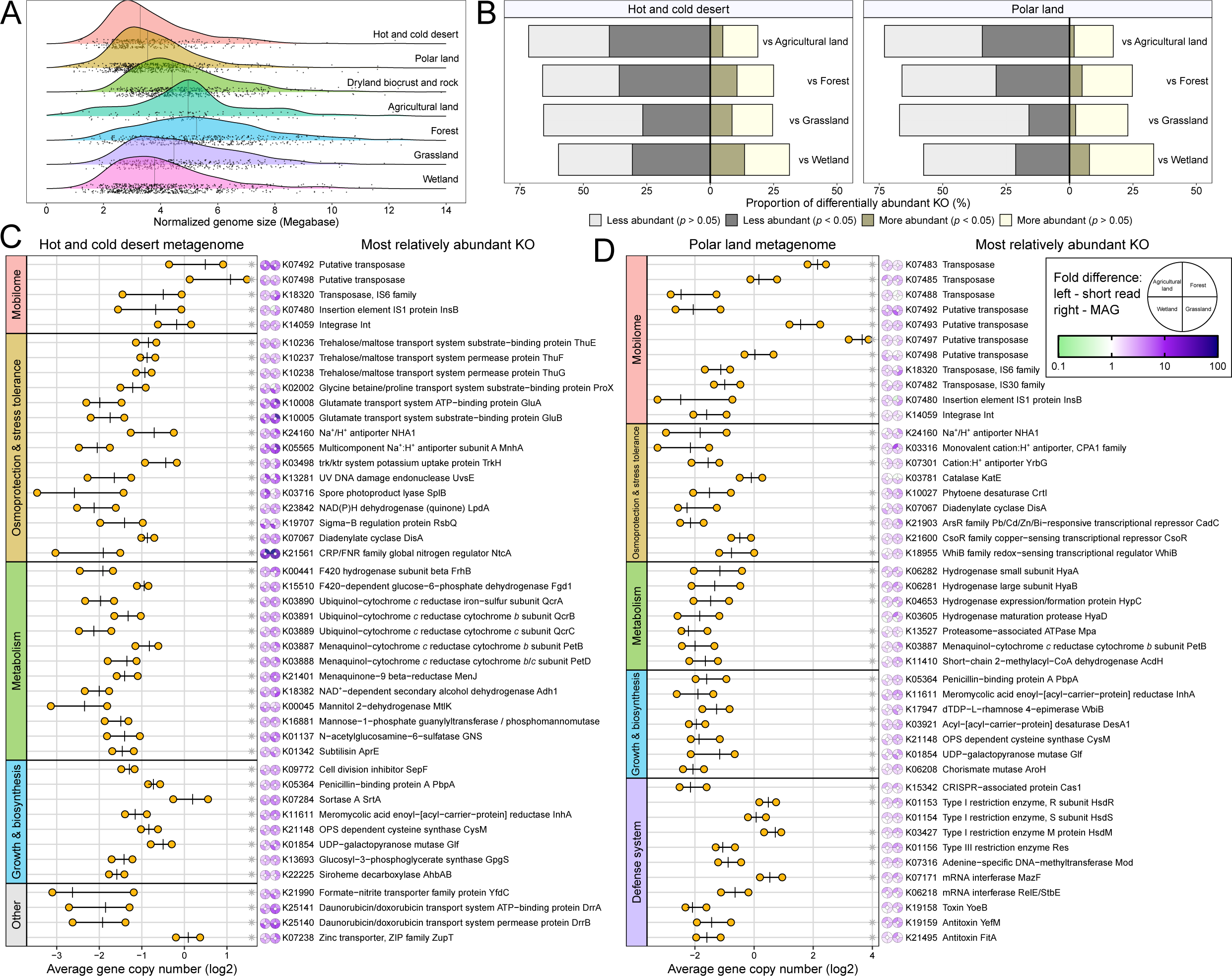
Functional signatures of the desert microbiome. (**A**) Distribution of completeness-normalized genome sizes of MAGs across different ecosystems. (**B**) Proportion of differentially abundant KEGG Orthology (KO) in nonpolar and polar desert microbiome compared to non-desert microbiome based on community gene copy numbers. Significance values were based on Welch’s t-test, corrected by Benjamini-Hochberg procedure. (**C-D**) The top 45 most relatively abundant KO (excluding uncharacterized proteins) with an average gene copy number over 0.25 in (**C**) hot and cold desert microbiomes and (**D**) polar desert microbiomes compared to non-desert soil microbiomes. The lollipop plot shows the lower quartile, median, and upper quartile of the average gene copy number of each KO in short read desert metagenomes. Asterisks denote the average gene copy number of the KO is more abundant in both nonpolar and polar desert communities compared to non-desert communities. The donut charts show the fold difference of each KO (left: short read data; right: MAGs) in nonpolar desert and polar land against other non-desert ecosystems.

### Desert microbes are enriched with mobile genetic elements and genes for antiviral defence, osmoregulation, and continuous energy harvesting

To gain a systematic overview of the desert microbial functions, we profiled the abundance and distribution of gene families defined by the KEGG protein database (KEGG Orthology; KO) (40) in both metagenomic short reads and derived MAGs (**Table S7**). Reflecting genome streamlining, 60 – 75% of gene families (defined by the KEGG protein database) have fewer average gene copies per genome in deserts compared to non-deserts (**Fig. 2B**). The reduction in genetic content appears to be primarily contributed by non-core functions. The functional categories most relatively depleted in nonpolar and polar deserts are membrane transport systems (e.g. ABC transporters and Major Facilitator Superfamily; av. 53 less gene copies/genome in desert communities), genes involved in energetically expensive processes (e.g. flagellar system for motility, antimicrobial resistance), protein kinases, and transcription factors, as well as structural proteins common in diderm bacteria (lipopolysaccharide biosynthesis, outer-membrane secretion systems) (**Fig. S3, Table S7**).

Unexpectedly, mobile genetic elements including numerous transposases and integrases were the most differentially enriched genes in desert ecosystems (**Fig. 2C-D**). Mobilome-associated gene families were 1.4 to 2.5-fold more prevalent in deserts, peaking in the hyper-arid Atacama Desert (av. 19,150 reads per million) (**Fig. S4A-B**). Given their reported roles (41–44), these mobile genetic elements may contribute to shape desert populations by 1) disseminating adaptive accessory traits by horizontal gene transfer, 2) increasing gene flow in otherwise slow-growing communities, and 3) promoting community turnover and nutrient release through viral lysis. In parallel, gene families involved in prokaryotic defence against foreign DNA are also highly enriched in desert communities (**Fig. 2C-D, Fig. S4C**), suggesting a regulatory process or a hidden arms race between the mobilome and the hosts. CRISPR-Cas, restriction-modification, and toxin-antitoxin systems are especially widespread among polar soils. In these bacteria-dominated desert ecosystems, the top-down pressure exerted by bacteriophages and mobile genetic elements may be a main overlooked driver of desert ecology.

Many of the most enriched gene families in deserts are involved in osmoregulation and stress tolerance, in support of previous reports (12, 19, 45). These include transporters of compatible solutes such as trehalose (ThuEFG), glycine betaine (ProX), and glutamate (GluAB), cation:H^+^ antiporters (MnhA, NHA1), and oxidative stress response proteins (UvsE, SplB, DisA, CrtI) (**Fig. 2C-D**). Some metabolism, growth, and biosynthesis genes were also highly enriched, though some probably reflect their conservation in monoderm bacteria ubiquitous in deserts rather than specific adaptations (e.g. menaquinone and cell wall synthesis proteins) (46–51) (**Fig. 2C-D**). Intriguingly, the top five most relatively abundant metabolism genes encode structural and maturation proteins for molecular hydrogen (H_2_) consumption (**Fig. 2C-D**) whereas the least relatively abundant KO are proteins involved in fermentation (e.g. Por, Pta, FhlA) and carbohydrate metabolism (e.g. XynA, GE, XylS) (**Fig. S3A-B**), inferring a unique metabolic lifestyle of microorganisms in these low carbon but highly oxygenated ecosystems.

### A shift from organoheterotrophy to lithoautotrophy in desert microorganisms

We next performed in-depth metagenomic profiling of a wide suite of genes to understand how desert microbes meet energy and carbon needs (**Table S6-8**). Desert microbes, though assumed to be predominantly organoheterotrophic (52), have reduced genomic capacity to degrade organic substrates compared to those found elsewhere. Based on gene-centric short read analysis, genes encoding carbohydrate-active enzymes (CAZy) glycoside hydrolases, polysaccharide lyases, and carbohydrate esterases were significantly less abundant in both nonpolar and polar deserts (all *p* < 0.005, Welch’s t-test) (**Fig. 3A**). Similarly, the two most abundant peptidase families (metallopeptidases and serine peptidases) were less abundant in deserts than agricultural, forest, and grassland soils (all *p* < 0.001, Welch’s t-test) (**Fig. 3B**). These inferences were also supported at the genome level (**Fig. 3A**), with consistent inferences made for the five most abundant desert bacterial orders (Solirubrobacterales, Acidimicrobiales, Mycobacteriales, Rubrobacterales, Gaiellales) (**Fig. 5C**) and after accounting for genome size (**Fig. S5**). Based on CAZy substrate preferences, desert microbes encoded fewer genes for using complex carbohydrates (cellulose, lignin, pectin), but were enriched with α-glucan-degrading enzymes (**Fig. 3C**). Likewise, enriched ABC transporters predominantly translocate simple sugars and α-glucosides in deserts (**Fig. S4D**). The copy number of various CAZy and peptidase families in dryland biocrust and rock-colonising (lithobiontic) communities is intermediate between desert soils and non-desert soils (**Fig. 3A-B**). Thus, low organic carbon input limits and shapes heterotrophic capabilities in desert soil communities.

**Figure 3.**
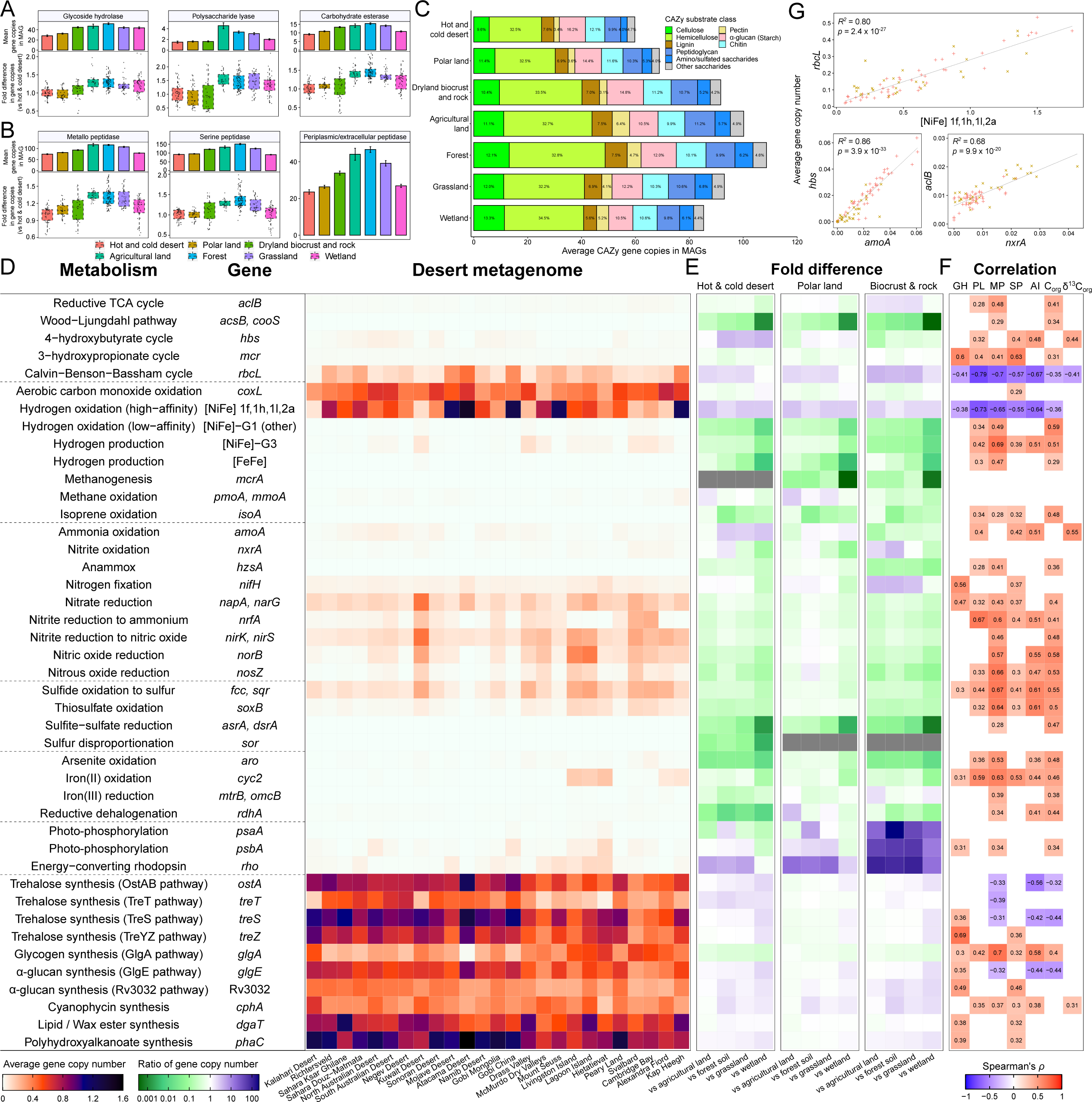
Metabolic capability of desert microbial communities versus non-desert ecosystems. (**A, B**) Boxplot showing the fold difference of community gene copy numbers of (**A**) carbohydrate active enzyme (CAZy) families and (**B**) peptidase families in various ecosystems against mean of hot and cold desert metagenomes based on gene-centric short read analysis. For boxplot, each data point is shown, and the horizontal lines indicate the 25th, 50th and 75th percentiles of the data, with vertical lines indicating the range, with outliers detached from the vertical lines. Bar charts show mean copy numbers of the genes in metagenome-assembled genomes (MAGs) recovered from the same environment. Periplasmic and extracellular peptidase implicated in the utilization of extracellular proteinaceous substrates encoded by MAGs were predicted by PSORTb. Copy numbers were normalized against completeness of each MAG (mean ± SEM). (**C**) Average copy numbers of CAZy targeting different substrate classes encoded by MAGs (normalized by completeness) from each ecosystem (mean ± SEM). Numbers within the bar chart denote the proportion of the preferred substrate class by CAZy within the same ecosystem. (**D**) Heatmap showing average copy numbers of metabolic marker genes involved in carbon fixation, trace gas cycling, nitrogen metabolism, sulfur metabolism, arsenite and iron oxidation, iron reduction, reductive dehalogenation, photophosphorylation, and energy reserve synthesis in desert communities. (**E**) Heatmap showing the fold difference of each metabolic marker gene in hot and cold desert, polar land, and dryland biocrust / rock against other non-desert ecosystems. (**F**) Spearman correlation of metabolic marker gene abundance with glycoside hydrolase (GH), polysaccharide lyase (PL), metallo-peptidase (MP), serine peptidase (SP), aridity index (AI), organic carbon content (C_org_), and δ^13^C signature of soil organic compound (δ^13^C_org_) in 75 samples. Correlations with aridity index are based on variables in nonpolar sites, as this metric is not accurately modelled for polar regions (66, 67). Only significant correlations with adjusted *p* values (Benjamini-Hochberg correction) smaller than 0.05 are shown. (**G**) Linear regressions of gene copy numbers of *rbcL* against high-affinity [NiFe]-hydrogenases, *hbs* against *amoA*, and *aclB* against *nxrA* in desert communities. *R^2^* and *p* values for each regression are shown.

The genomic capacity for autotrophy showed the opposite trend to heterotrophy. Considering both chemosynthetic and photosynthetic carbon fixation pathways, 21% of desert microbes are primary producers. RuBisCO genes (*rbcL*) were among the most abundant metabolic genes in deserts (17% cells; **Fig. 3D**), and were enriched by two-fold in deserts compared to non-desert sites (except for wetlands where there is much anaerobic chemolithoautotrophy (53); **Fig. 3E**). Consistently, stable isotope signatures suggest desert organic matter (δ^13^C_org_) primarily originates from RuBisCO-based carbon fixation (54, 55), with ^13^C depletion within the range of -19.5‰ in North Australian Desert to -28.2‰ in the Atacama Desert (**Fig. 6C**). While plant RuBisCO likely contributes to this signature, microbial RuBisCO likely predominates at hyper-arid sites; consistently, the abundance of *rbcL* increases with decreasing soil organic carbon (*ρ* = -0.35, *p* < 0.01) and aridity (*ρ* = -0.70, *p* < 0.001), with similar observations for the δ^13^C_org_ values, whereas CAZy and peptidases follow opposite trends (**Fig. 3F & Table S9**). The archaeal 4-hydroxybutyrate cycle is also selectively enriched in deserts (**Fig. 3D-E**). Primary producer abundance was also elevated in dryland biocrusts and rocks (36%) where photoautotrophic and diazotrophic Cyanobacteria are ubiquitous (**Fig. 3E, Fig. S1**). Together, these findings suggest the physicochemical conditions and resultant low vegetation-derived carbon inputs in deserts select for metabolically self-sufficient autotrophs.

Autotrophy was strongly associated with aerotrophy and nitrification, not photosynthesis, in non-biocrust desert soils (**Supplementary Note 2**). RuBisCO abundance was most strongly correlated with high-affinity [NiFe]-hydrogenases (*R^2^* = 0.80; **Fig. 3G**), with genes encoding both enzymes enriched in deserts overall and highest in the hyper-arid Atacama (the site with the strongest ^13^C_org_ depletion; **Fig. 6C**). These inferences were also supported by MAG-level analyses (**Fig. S6B**), with two thirds of RuBisCO-encoding MAGs co-encoding uptake hydrogenases, including multiple Actinobacteriota, Chloroflexota, and Dormibacteria orders (**Fig. 5D, Table S6**). In further evidence of the dominant role of aerotrophy, nearly half of desert microbes encode enzymes to consume carbon monoxide (some alongside RuBisCO; **Fig. 3D**, **Fig. 5D**), while methane oxidation was an enriched rarer trait primarily mediated by the abundant atmospheric methanotroph *Ca.* Methyloligotrophales (USCγ (21, 56); **Fig. 3E**, **Fig. 5B**). Nitrification also appears to be a key primary production process: based on gene correlations and MAG-level analyses, the 4-hydroxybutyrate and reductive tricarboxylic cycles were strongly associated with archaeal nitrification (Nitrososphaerales) and bacterial nitrification (Nitrospirales), respectively (**Fig. 3C**, **Fig. 5D**), with some nitrifiers also likely persisting on H_2_ as per culture-based inferences (57, 58) (**Fig. 5D**). Numerous bacteria encode genes to harness other lithic substrates, including sulfide (6.1% cells), thiosulfate (4.2%), ferrous iron (2.7%), and arsenite (0.70%) (**Fig. 3D**, **Fig. 5D**).

Our analyses suggest continuous energy harvesting and energy reserve storage are complementary strategies that are both enriched in desert ecosystems. While most desert microbes can continually persist on atmospheric H_2_ and CO, some eight bacterial orders (e.g. Rubrobacterales) also encoded energy-converting rhodopsins to harvest sunlight and these genes were especially enriched in biocrust and rock-associated communities (**Fig. 3D**, **Fig. 5D; Supplementary Note 2**). While there was minimal capacity for oxygenic photosynthesis, a small number of Proteobacteria encoded genes to harvest sunlight via photosystem II (**Fig. 5D**). Energy reserve metabolism is an extremely widespread trait. The capacity to synthesize trehalose (present in 76 orders; also an osmoprotectant), glycogen (61 orders), glycogen-like α-glucans (81 orders), cyanophycin (39 orders), triacylglycerol / wax ester (32 orders), and polyhydroxyalkanoate (53 orders) are ubiquitous among desert MAGs (**Fig. 5D**). These storage compound biosynthesis genes were most abundant in nonpolar deserts compared to non-deserts (**Fig. 3E**). Remarkably, the Atacama Desert exhibited the strongest potential for trehalose, α-glucan, and polyhydroxyalkanoate synthesis, while cyanophycin is comparatively enriched in polar lands. These results lend support to the hypothesis that energy reserve metabolism enables adaptation to the pulse dynamics in hot and cold deserts, with microbes rapidly accumulating energy reserves during brief “metabolic windows” after precipitation to prepare for extended periods of energy scarcity (4, 59).

### Aridity as the primary driver of functional microbiome and edaphic conditions

We applied both random forest and Spearman correlation analyses to determine the drivers underlying the trends of microbial and gene abundance observed in the above comparative analysis (**Fig. 4**). The importance of eight representative non-collinear (|*ρ*| < 0.7) climatological predictors, including aridity, annual mean temperature, seasonality of precipitation and temperature, mean diurnal temperature range, wind speed, soil C:N ratio, and elevation of both desert and non-desert ecosystems (n = 275), was assessed. Random forest and correlation analysis both suggested aridity is the best predictor, by a large margin, of the abundance of microbial taxa. Similarly, a disproportionate fraction of KO, CAZy, and metabolic marker genes are best predicted by aridity (46 – 59% of genes), followed by precipitation seasonality (4 – 14%) and annual mean temperature (4 – 10%). Abundance of these genes are also most highly correlated with aridity (av. |*ρ*| = 0.30 for KO, 0.34 for CAZy, 0.38 for metabolic markers), more so than any other predictors (**Fig. 4, Fig. S7E-F**). Most of KO and CAZy are significantly positively correlated with aridity (**Fig. S7A,C**), in line with the observation that a large fraction of genes in arid desert communities have fewer copies per genome (**Fig. 2B**). Correspondingly, the majority of the most relatively more abundant KO and less abundant KO in desert communities (**Fig. 2C-D**, **Fig. S3A-B**) are also significantly negatively and positively correlated with aridity, respectively (**Fig. 4A**), whereas few are significantly correlated with annual mean temperature (**Fig. S7B**).

**Figure 4.**
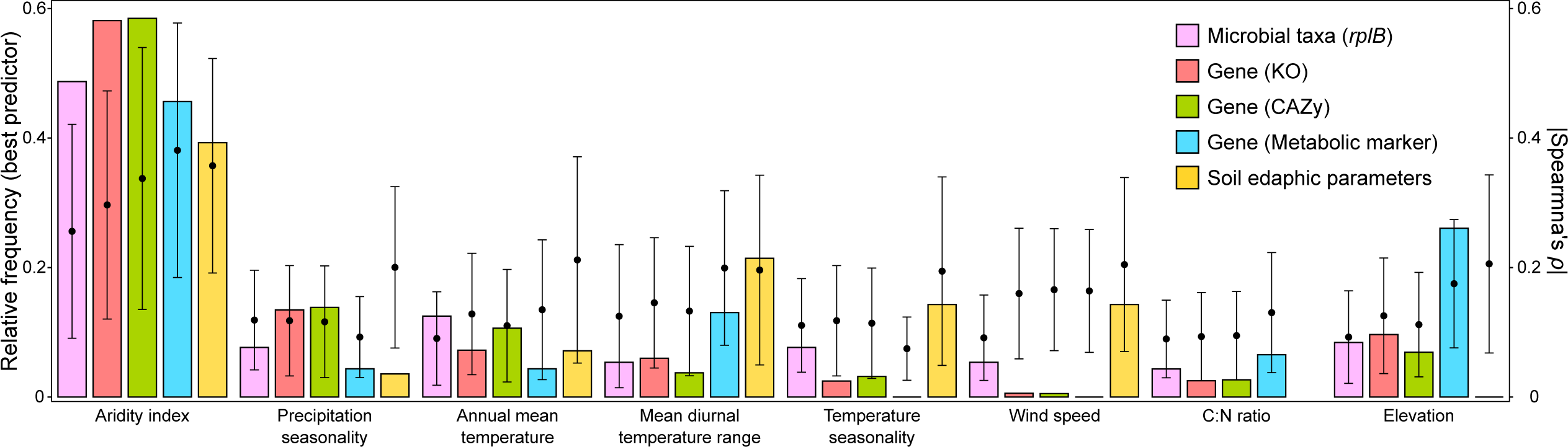
Aridity as the primary climatological driver of microbial abundance, gene abundance, and edaphic conditions in soil ecosystems. Bar chart showing relative frequency of the eight representative climatological parameters determined by random forest analysis as the best predictor for abundance of microbial genera, KEGG Orthology (KO), carbohydrate-active enzymes (CAZy), and metabolic maker genes in soil ecosystems (n = 275) and measured edaphic parameters of deserts (n = 75). Mean of absolute value of Spearman correlation of gene abundance and edaphic parameters with each climatological parameter was also shown (mean ± sd).

The effect of aridity on microbial functions is likely mediated by modulation of organic and litter input (e.g. *ρ* of aridity and Normalized Difference Vegetation Index = 0.86), water availability, and other soil physicochemical conditions. To evaluate this, we performed the same driver analyses for edaphic parameters measured in sampled desert soils. Consistent with gene abundance, edaphic conditions are most significantly modulated by aridity, as suggested by both random forest and correlation analyses (**Fig. 4**), including organic carbon (*ρ =* 0.44), electrical conductivity (*ρ =* -0.57), and manganese (*ρ =* 0.64) (all *p* < 0.05) (**Table S9**). Together, these results suggest that global scale gene abundance patterns and adaptations of soil communities are primarily controlled by ecosystem water availability.

### Desert microorganisms mediate respiration, trace gas oxidation, chemosynthesis, and photosynthesis at rates correlated with aridity

Metagenomic analysis suggests that a combination of organic, inorganic, and solar energy sources sustain the energy and carbon needs of desert communities. We evaluated these inferences by profiling activities and potentials of multiple metabolic processes in soils from each site. Independent whole soil microcosms were set up to measure: (i) soil CO_2_ emission as a proxy for community respiration and organotrophic activities; (ii) oxidation of H_2_, CO, and CH_4_; and (iii) fixation of radiolabelled ^14^CO_2_ in dark, H_2_-stimulated, and illuminated conditions. For dry soils at their native soil water content, respiration was detected in 13 of the 25 sites, mostly from polar locations. In contrast, following rehydration, all soils had measurable CO_2_ emissions. Soil respiration rates range from 0.01 to 18 µmol g_dw_^-1^ day^-1^, with the Atacama Desert (among nonpolar deserts) and McMurdo Dry Valleys (among polar deserts) soils exhibiting the lowest, albeit still quantifiable, respiration rates (**Fig. 6A**). Thus, viable and active populations reside within these environments. Respiration rates were best predicted by total cell counts (*ρ* = 0.71, *p* < 10^-10^) and organic carbon (*ρ* = 0.50, *p* < 10^-4^) (**Fig. 6E**), positively correlated with the abundance of CAZy and peptidases, and negatively correlated with relative abundance of RuBisCO and hydrogenases (**Table S9**).

**Figure 5.**
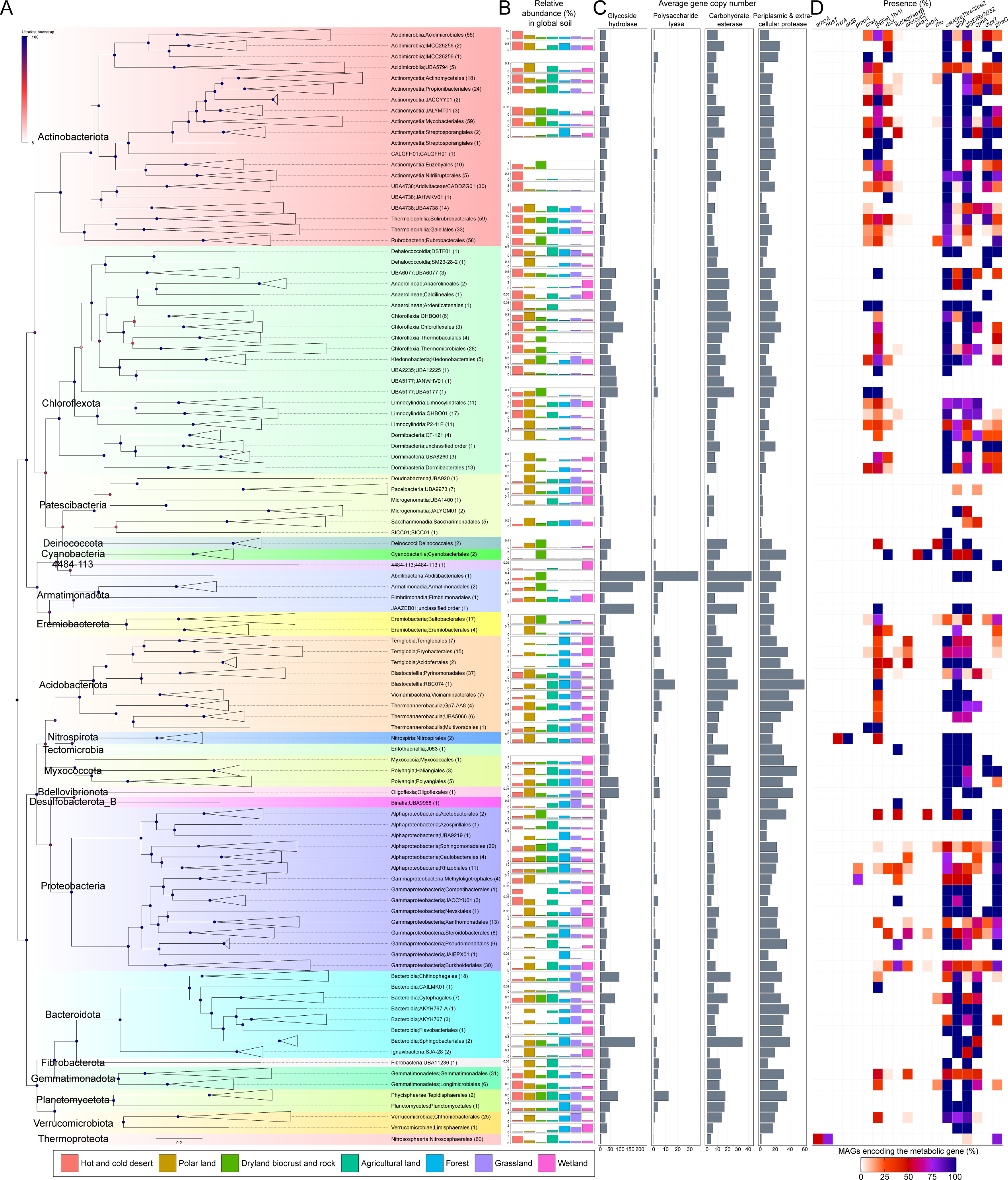
Phylogenetic diversity, genome size distribution, abundance, and metabolic capability of 911 medium and high-quality metagenome-assembled genomes (MAGs) from global deserts. (**A**) Maximum-likelihood phylogenetic trees of 851 bacterial MAGs based on 120 GTDB bacterial marker genes with leaves collapsed at the order level (total 102 orders). All 60 archaeal MAGs were from a single order (Nitrososphaerales) and are presented below the bacterial tree. Tip labels indicate class and order level GTDB release 09-RS220 taxonomy and brackets show number of MAGs from the order. Nodes are colored based on branch support by 1000 ultrafast bootstrap replicates. (**B**) Mean relative abundance of the order affiliated with desert MAGs across different ecosystems based on the ribosomal marker *rplB*. (**C**) Mean copy numbers of degradative glycoside hydrolase, polysaccharide lyase, carbohydrate esterase, and periplasmic and extracellular peptidases in MAGs of each order. The value was normalized against MAG completeness. (**D**) Prevalence of metabolic marker genes involved in inorganic compound oxidation, photophosphorylation, carbon fixation, and energy reserve synthesis (surveyed in Fig. 3D) in MAGs of each order.

**Figure 6.**
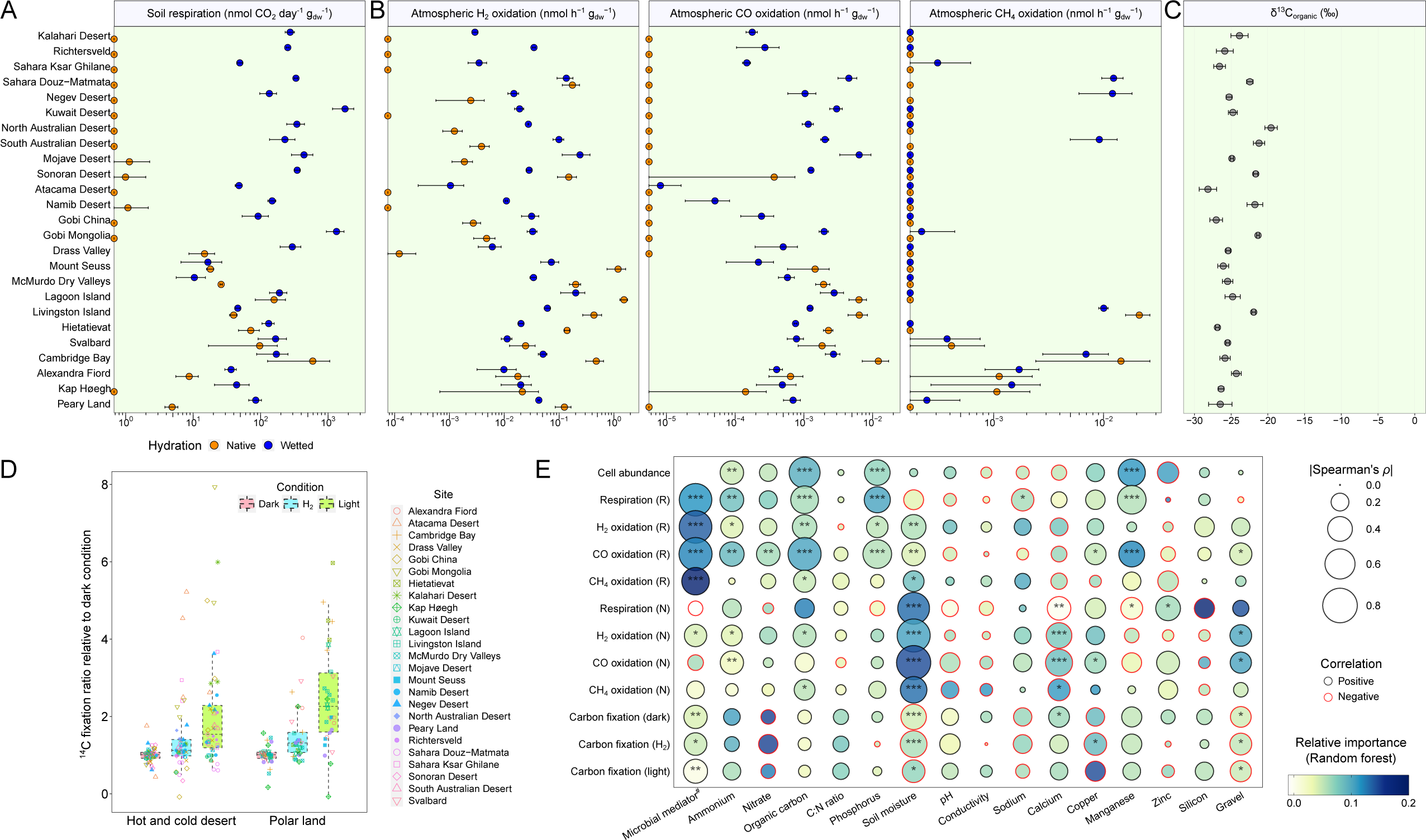
Activity and drivers of respiration, trace gas oxidation, ^13^C isotopic signature, and ^14^CO_2_ fixation of desert soils. (**A**) Soil respiration (CO_2_ emission) and (**B**) atmospheric H_2_, CO, and CH_4_ oxidation (mean ± standard error of biological triplicates) at native and wetted hydration conditions during microcosm incubations (20°C for hot and cold desert samples; 10°C for polar lands and Drass Valley). Rates were normalized to dry weight and corrected against heat-killed controls. (**C**) Depletion of ^13^C isotope in soil organic matter. (**D**) Relative rates of ^14^CO_2_ fixation for pooled hydrated soils in technical triplicate under dark ambient, hydrogenotrophic (100 ppmv of headspace H_2_), and photosynthetic conditions (40 μmol photons m^-2^ s^-1^). (**E**) Random forest driver analysis and Spearman correlation of 12 microbial activity parameters (cell abundance, respiration, H_2_, CO, and CH_4_ oxidation, dark, hydrogenotrophic, and photosynthetic carbon fixation) with 16 representative edaphic variables. “R” and “N” denote rehydrated and native hydration conditions, respectively. Microbial mediator refers to abundance of all cells (respiration), H_2_ oxidizers (H_2_ oxidation), CO oxidizers (CO oxidation), CH_4_ oxidizers (CH_4_ oxidation), all autotrophs (dark C fixation), CBB-autotrophs (hydrogenotrophic C fixation), and photoautotrophs (photosynthetic C fixation). Other edaphic variables were selected to minimize multicollinearity (Spearman’s |ρ| < 0.7 in all cases). Significant Spearman correlation is denoted by asterisks (adjusted *p* value ≤ 0.001, ***; ≤0.01, **; ≤0.05, *).

Activity measurements confirmed atmospheric H_2_ and CO are major energy sources for desert communities. All desert soil communities consumed both gases under rehydrated conditions (**Fig. S8**), with average rates greatly varying between sites, from 0.001 to 0.24 nmol H_2_ g_dw_^-1^ hr^-1^ and 0.01 to 6.4 CO pmol g_dw_^-1^ hr^-1^ (**Fig. 6B**). In line with the variable and relatively low abundance of methanotrophs, consumption of CH_4_ only occurred at 11 sites, at rates between 0.25 to 12 pmol g_dw_^-1^ hr^-1^ (**Fig. 6B**). These biogeochemical and metagenomic findings support previous observations that desert soils are quantitatively significant sinks of atmospheric CH_4_ (60). Consumption rates of H_2_, CO, and CH_4_ in wetted soils were best predicted by the abundance of H_2_ oxidizers (*ρ* = 0.71), CO oxidizers (*ρ* = 0.73), and CH_4_ oxidizers (*ρ* = 0.50), respectively; these activities also significantly correlate with organic carbon levels (**Fig. 6E**). H_2_ and CH_4_ uptake activities do not significantly correlate with respiration, whereas CO oxidation does (*ρ* = 0.48) (**Table S9**). We also tested whether desert communities were capable of continuously harvesting trace gases under native dry conditions (**Table S1**). Atmospheric H_2_, CO, and CH_4_ were consumed by soils from 19, 10, and five sites respectively (**Fig. 6B**). For polar sites, trace gas oxidation rates were often higher at native hydration, potentially because gases more readily diffuse through native unsaturated soils (61, 62). Under native hydration, oxidation rates of H_2_ (*ρ* = 0.73), CO (*ρ* = 0.78), and CH_4_ (*ρ* = 0.46), as well as respiration (*ρ* = 0.67), are governed by soil moisture rather than cell abundance (**Fig. 6E**). Altogether, these results confirm trace gases, especially H_2_, are key energy sources for desert communities and enable some energy harvesting even under dry conditions.

All sampled soils also mediated chemosynthesis and photosynthesis. Across the three conditions tested (dark, light, H_2_-stimulated), ^14^CO_2_ assimilation rates commonly fall within 0.01 to 1 nmol ^14^CO_2_ g_dw_^-1^ day^-1^ (**Fig. S9**), though gross fixation rates are likely much higher given soil inorganic carbon invariably competes with the labelled ^14^CO_2_ (63). Dark CO_2_ incorporation was measured at all sites, likely reflecting anaplerotic assimilation of CO_2_, given the absolute rate correlates most significantly with respiration (*ρ* = 0.58) (**Table S9**). Yet, dark CO_2_ assimilation also correlates with abundance of soil autotrophs (*ρ* = 0.35) (**Fig. 6D**), suggesting possible chemosynthesis supported by the presence of inorganic edaphic substrates such as ammonium in soils (**Fig. S1B**). Confirming metagenomic predictions that desert microbes coupled H_2_ oxidation to carbon fixation, ^14^CO_2_ fixation significantly increased by 40% in nonpolar deserts and 52% in polar deserts under H_2_-stimulated conditions (all *p* < 0.01, Welch’s t-test). Despite low initial abundance of photoautotrophs, light stimulation increased ^14^CO_2_ fixation by 2.1- and 2.5-fold respectively (**Fig. 6D**); this likely reflects that photoautotrophs rapidly responded and proliferated in the presence of water and constant illumination (64, 65) during the five-day incubation. The dominant CO_2_ fixation processes varied between the sites, with H_2_ most strongly stimulating fixation in the Atacama, Svalbard, and Alexandra Fiord samples, whereas light had the strongest effect in the Gobi Mongolia, Kalahari, and Hietatievat sites. Corroborating metagenomic inferences, these results confirm a widespread capacity for microbial primary production and an overlooked contribution of chemosynthesis in desert soils.

## Conclusion

Our study demonstrates that common biological constraints across geographically distant deserts select for compositionally similar microbial communities with conserved metabolic adaptations. Much of the variation in microbiome structure, function and activity was driven by aridity, either directly, or through its effects on soil characteristics.

Several capabilities define microbial communities in deserts compared to other terrestrial environments. In addition to osmoprotection and stress tolerance, enigmatic mobile genetic elements and prokaryotic defence systems are highly enriched, suggesting unknown mechanisms of evolution and adaptation of oligotrophic desert microorganisms. With limited carbon input from vegetation in desert soils, a consistent trend is an increased reliance on lithoautotrophy over organoheterotrophy. Observations of reduced copy numbers of CAZy and peptidases for degradation of carbohydrates and proteins, together with wider genome streamlining, provide support for this idea. In contrast, facultatively hydrogenotrophic autotrophs represented by diverse members of Actinobacteriota and Chloroflexota increase in abundance with increasing aridity and decreasing organic carbon available. Ammonia-oxidizing Nitrososphaerales and nitrite-oxidizing Nitrospirales are also ubiquitous autotrophs in deserts. Collectively, they contribute significantly to chemosynthetic activities in these soils, together with photoautotrophs under favourable conditions. Under drought, rates of these biogeochemical activities are dictated by moisture levels, whereas abundance of corresponding microbial mediators are the best predictors at rehydration. Our results suggest that most desert microbes have the capacity both to continuously harvest energy, using atmospheric trace gases and sometimes sunlight (primarily via rhodopsins), but also to synthesize and metabolise energy reserves. The latter appears to be a particularly relevant strategy in nonpolar deserts where communities often experience pulse hydration dynamics.

By evaluating trends across aridity, our research provides critical insights to predict potential shifts of the structure and function of soil microbiome amid global desertification. We predict aridification will lead to a simplification in soil microbial community and functional diversity. This will be accompanied by a reduction in photoautotrophic and heterotrophic capacity, only partially offset by an increased contribution from chemolithoautotrophy. Nevertheless, overall primary productivity will likely decrease with a smaller carrying capacity than non-desert environments and thereby slow overall elemental cycling. However, increased moisture driven by glacial melt and permafrost thawing in warming polar lands and shifted hydrology in some nonpolar deserts may drive opposite trends. Interdisciplinary research with reciprocal experiments is needed to resolve these dynamics and their broader consequences as climate change and human activity continue to accelerate.

## Materials and Methods

### Field sampling and site characterization

We collected triplicate soil samples from 25 locations spanning hot, cold, and polar desert landscapes across 15 countries and all seven continents, including ten hot desert sites (Kalahari Desert, South Africa; Richtersveld, South Africa; Ksar Ghilane region of Sahara Desert, Tunisia; Douz-Matmata region of Sahara Desert, Tunisia; Kuwait Desert, Kuwait; Negev Desert, Israel; North Australian Desert, Australia; South Australian Desert, Australia; Sonoran Desert, USA; Mojave Desert, USA), five cold desert sites (Namib Desert, Namibia; Mongolia region of Gobi Desert, Mongolia; China region of Gobi Desert, China; Drass Valley, India; Atacama Desert, Chile), and ten polar sites (McMurdo Dry Valleys, Antarctica; Mount Seuss, Antarctica; Livingston Island, Antarctica; Lagoon Island, Antarctica; Hietatievat, Finland; Svalbard, Norway; Cambridge Bay, Canada; Alexandra Fiord, Canada; Peary Land, Greenland; Kap Høegh, Greenland). The surveyed hot and cold desert sites are characterized by an aridity index below 0.2 and classified as Deserts and semi-deserts biome (T5) within the IUCN Global Ecosystem Typology (2) (except for the montane Drass Valley). Polar desert sites are characterized by an annual mean temperature below 0°C (1) and classified as polar tundra and deserts (T6.3) or polar/alpine cliffs, screes, outcrops and lava flows (T6.2) within the IUCN Global Ecosystem Typology (2) (except for the inland sand dune Hietatievat). The sampled locations are respectively classified as and Polar tundra and deserts (T6.3) within. As detailed in **Table S1**, sampling in most locations was performed during dry seasons between 2013 and 2019 and on open areas with minimal vegetation cover and plant materials. At each site, the upper 10 cm of soils were collected aseptically from three different spots of at least 50 m apart, i.e. biological triplicate per sampling location. After collection, mineral soils from polar and cold mountainous sites were immediately stored at -20°C, while other dry desert samples were stored at ambient temperatures. Samples were subsequently transported to Monash University’s quarantine approved facilities for standardized laboratory analyses.

Climatological and geospatial information of sampling sites were obtained from the WorldClim database v2.1 (mean annual precipitation, mean annual temperature, wind speed, elevation, solar radiation, water vapour pressures, and 19 BioClimate variables) (66), Global Aridity Index and Potential Evapo-Transpiration (ET0) Database v5 (aridity index and potential evapotranspiration) (67), and NASA’s Oak Ridge National Laboratory Distributed Active Archive Center (C:N ratio of soil microbial biomass (30 cm) (68), leaf area index (LAI) (69), normalized difference vegetation index (NDVI) (70), and gross primary productivity (GPP) (71)). Aridity index was retrieved for nonpolar sites only as the matric is not accurately modelled for polar regions (66, 67). A principal component analysis on z-score normalized climate variables was performed by base R function *pca* and visualized using the biplot function in PCAtools v2.8.0 (https://github.com/kevinblighe/PCAtools). Detailed information on individual sample including GPS coordinates, elevation, and other metadata can be found in **Fig. S1** and **Table S1**.

### Soil physicochemical characterization

To characterize soil physicochemical parameters, an equivalent amount of soil from each of the three replicates of the same site was first aliquoted and pooled as a composite sample. Each sample was treated with gamma irradiation at an absorbed dose of 50 kGy (Steritech Pty Ltd Victoria, Australia) for compliance with Australian Department of Agriculture, Water and the Environment’s quarantine good regulations. They were subsequently transported to the Environmental Analysis Laboratory (EAL), Southern Cross University, Australia for physicochemical assays in accordance with ISO/IEC 17025 standard procedures. A comprehensive list of physicochemical parameters were analysed as follows: basic soil color and texture; grain size; pH and electrical conductivity (1:5 water); total carbon, nitrogen, organic carbon, and organic matter; available calcium, magnesium, potassium, ammonium, nitrate, phosphate, sulfur; exchangeable sodium, potassium, calcium, magnesium, hydrogen, and aluminium; effective cation exchange capacity; Bray I, Bray II, and Colwell phosphorus; and available micronutrients zinc, manganese, iron, copper, boron, and silicon. Measurement data and corresponding analytical methods are provided in **Table S1**. We further characterized soil gravimetric water content for each individual soil in house. Approximately 2 - 3 g of soil, in technical duplicate, was weighed on a clean aluminium weigh tin using a precision balance (± 0.001 g) and dried at 105°C for 24 h. Prior to re-weighing, the sample was cooled within a desiccator to minimize absorption of air moisture. The sample weight was recorded and dried again at 105°C for 24 h. The procedure was repeated until no further change in weight was observed. Gravimetric water content was calculated by dividing the weight difference with the original sample weight.

### Soil DNA extraction and quantitative PCR

Community DNA for metagenomic sequencing was extracted from 0.5 g of soil using the FastDNA™ SPIN Kit for soil (MP Biomedicals) according to the manufacturer’s instructions and eluted with 30 µl UltraPure™ DNase/RNase-free distilled water (Invitrogen), except for the very low biomass Atacama soils of which repeated extractions yielded no quantifiable DNA. We therefore extracted DNA from Atacama samples using a modified CTAB extraction method as described in Warren-Rhodes et al. 2019 (10). 0.5 g of soil was combined with 0.25 g each of 0.1 mm and 0.5 mm silica-zirconia beads, and 320 µl each of 100 mM NaH_2_PO_4_ buffer and SDS lysis buffer (100 mM NaCl, 500 mM Tris pH 8.0, 10% SDS) in a close-top screw tube. The sample was disturbed with the FastPrep®-24 (MP Biomedicals) at speed 5.5 for four rounds of 30 s beating and 30 s of ice cooling, which was then centrifuged at 13,200 rpm for 3 min. Following addition of 230 μl of hexadecyltrimethylammonium bromide (CTAB) extraction buffer (100 mM Tris-HCl, 1.4 M NaCl, 20 mM EDTA, 2% CTAB, 1% polyvinylpyrrolidone-MW360000 and 0.4% 2-mercaptoethanol), samples were homogenised by 10 s vortex, incubated at 60°C and 300 rpm for 30 min, and then centrifuged at 13200 rpm for 1 min. Each sample was added with 550 μl of chloroform:isoamyl alcohol (24:1), vortexed for 10 s and centrifuged at 13200 rpm for 5 min. The upper aqueous layer was transferred into an Eppendorf tube and amended with 10 M ammonium acetate to a final concentration of 2.5 M. The content was inverted and centrifuged at 13200 rpm for 5 min to further remove debris. The aqueous phase was transferred into a new Eppendorf tube and mixed with 0.54 volume of ice-cold isopropanol through inversion. The sample was then incubated at -20°C for 24 h for DNA precipitation. The content was centrifuged at 4°C and 13200 rpm for 60 min. The supernatant was carefully discarded, and the pellet was washed with 1 ml ice-chilled 70% ethanol and centrifuged at 4°C and 13200 rpm for 30 min. The supernatant was decanted, and the pellet was allowed to air dry. Replicate extractions of the same sample were resuspended and combined in 20 µl UltraPure™ DNase/RNase-free distilled water. Two blank extraction controls for FastDNA kit and one blank extraction control for CTAB method were also prepared. DNA extracts immediately stored at -20°C until further experiment. The quality and quantity of DNA were tested using a NanoDrop® ND-1000 spectrophotometer (Thermo Fisher Scientific) and a Qubit 2 fluorometer with dsDNA HS Assay Kit (Thermo Fisher Scientific), respectively.

Quantitative polymerase chain reaction (qPCR) assays were carried out to quantify the copy numbers of 16S rRNA genes for estimation of the total bacterial and archaeal cell count in each soil. The V4 hypervariable region of the 16S rRNA gene of the extracted DNA was amplified using the revised universal Earth Microbiome Project primer pairs F515 (Parada) (72) and R806 (Apprill) (73). PCR reactions were set up in a 96-well plate with each well containing 1 × 480 SYBR Green I Master Mix (Roche), 0.2 μM of each primer, and 1 μl of 1:100 diluted DNA and ultrapure water in a final volume of 20 μl on a LightCycler 480 real-time PCR system (Roche, Basel, CH). For each run, triplicate DNA samples were analysed and duplicate serial dilutions (10^1^ - 10^8^ copies) of DNA from a synthetic pUC-like cloning vector containing *E. coli* 16S rRNA gene sequence (pMA plasmid; GeneArt, Thermo Fisher Scientific) were included as standards for gene copy calibration (average crossing point values, C_p_, against gene copies). The qPCR conditions were as follows: pre-incubation at 95 °C for 3 min and 50 cycles of denaturation at 95 °C for 30 s, annealing at 54 °C for 30 s, and extension at 72 °C for 24 s.

### Metagenomic sequencing, assembly and binning

DNA was shipped on dry ice to the Australian Centre for Ecogenomics, University of Queensland. Metagenomic shotgun libraries were prepared using the Nextera XT DNA Library Preparation Kit (Illumina) and subject to paired-end sequencing (2 × 150 bp) on a NextSeq500 platform (Illumina). On average, sequencing yielded 31,400,000 read pairs for each of the 75 soil metagenomes and 34,870 read pairs for the three negative controls (**Table S2**), indicating minimal contamination from DNA extraction and sequencing processes. Raw metagenomic reads were quality processed through the BBDuk function of BBTools v38.68 (https://sourceforge.net/projects/bbmap/). The rightmost base known to contain a high error rate, sequencing adapters (k-mer size of 23 and hamming distance of 1), PhiX spike-in (k-mer size of 31 and hamming distance of 1), bases from 3’ ends with a Phred score below 20, and resultant reads with lengths shorter than 50 were sequentially removed. Read quality before and after processing was assessed using FastQC v0.11.7 (http://www.bioinformatics.babraham.ac.uk/projects/fastqc/). 91% high-quality read pairs were retained for downstream analysis.

Individual metagenomic assembly and co-assembly of the triplicate samples originating from the same location were performed with metaSPAdes v3.14.0 (74) and MEGAHIT v1.2.9 (75) (min k: 27, max k: 127, k step: 10), respectively. Quality processed short reads were mapped to the assembled contigs using Bowtie2 v2.3.5 (76) with default parameters to generate contig coverage profiles. Binning was performed using Autometa v1.0.2 (77), CONCOCT v1.1.0 (78), MaxBin2 v2.2.7 (79), and MetaBAT2 v2.15 (80) on contigs with length over 2000 bp. Resulting bins from the same assembly were then dereplicated using DAS_Tool v1.1.2 (81) and refined using RefineM v0.1.1 (82) to filter spurious contigs with incongruent genomic and taxonomic properties. Applying a threshold average nucleotide identity of 99%, bins from different assemblies were consolidated to a non-redundant set of metagenome-assembled genomes (MAGs) using dRep v3.0.0 (83). Only MAGs with completeness > 50% and contamination < 10%, as assessed by CheckM v1.1.3 (84), were retained. In total, 136 high quality (completeness > 90% and contamination < 5%) and 775 medium quality (completeness > 50% and contamination < 10%) (85) were recovered. Their taxonomy was assigned by GTDB-Tk v2.4.0 (86) with reference to the Genome Taxonomy Database (GTDB) R09-RS220 (87). Based on GTDB relative evolutionary divergence (RED) and average nucleotide identity (ANI) values, the 911 MAGs constituted an 814 candidate species (at 95% ANI) (**Table S6**).

### Retrieval and analysis of high-quality published metagenomes from global soil ecosystems

A total of 200 published metagenomes from non-desert ecosystems (spanning forest (n = 75) (30, 88–99), grassland (n = 33) (89, 100–108), agricultural land (n = 24) (109–115), and wetland (n = 42) (53, 104, 116–125)) and biocrust and lithic niches of drylands (n = 26) (27, 107, 126–130) were retrieved (**Table S3**). To ensure comparability and quality, public soil data was selected based on a compatible sampling and sequencing scheme with our current study: (i) surface soils were collected at depths between 0 – 30 cm in biological triplicates; (ii) sequenced natural soils were unamended; (iii) GPS coordinates of the sites were documented; (iv) community DNA was sequenced on an Illumina platform with a minimum sequencing depth of one million reads and a read length output longer than 130 bp; (v) metagenomes were from published studies and (vi) made publicly available; and (vii) overall sample set captured a broad geographic representation. Quality filtering, assembly and binning of the metagenomes were performed as described above, except that all assemblies were done using Megahit v1.2.9 (75) and that Autometa v1.0.2 (77) was not used for binning. A total of 2226 medium and high-quality MAGs from 45 phyla were recovered. Climatological and geospatial information of the sites were obtained as described above.

### Community analysis

Soil microbial community structures were determined independently by using both universal single copy ribosomal marker genes and small subunit rRNA (SSU rRNA) gene in metagenomes. SingleM v.0.13.2 (131) was used to map quality-filtered reads to fixed windows of conserved single copy ribosomal marker genes and the matched sequences were clustered to operational taxonomic units (OTUs) at 97% identity. Taxonomy to mapped reads was assigned using GTDB, allowing taxonomic consistency between community and MAG analysis. Reads assigned to eukaryotes were filtered. For community profiling, *rplB* was used as it has been benchmarked to be a robust taxonomic marker for delineating closely and distantly related bacteria and archaea while minimizing bias from copy number variations prevalent in 16S rRNA gene (132). PhyloFlash v.3.4 was used to screen quality-filtered reads for the small subunit ribosomal RNA sequences and assembled with the option –almosteverything (133). The SSU Ref NR99 database from the SILVA release 138.1 (134) served as the reference for the sequence searching and taxonomy assignment of SSU reads to the Nearest Taxonomic Units (NTUs). Full lists of OTUs and NTUs determined by both methods can be found in **Table S4 and S5**.

Differentially abundant taxa in nonpolar desert and polar sites compared to non-desert environments were identified using a combination of four independent analyses (ALDEx2 v1.30.0 (135), ANCOMBC v2.0.2 (136), LinDA v1.1 (137), MaAsLin2 v1.12.0 (138)) as per recent recommendation (139). Each tool was run on non-rarefied *rplB* datasets in default parameters with slight modifications: ALDEx2 (mc.samples = 1000), ANCOMBC (struc_zero = TRUE, conserve = TRUE), LinDA (prev.filter = 0.1, zero.handling = ’pseudo-count’, pseudo.cnt = 0.01), and MaAsLin2 (standardize = FALSE). All *p* values were adjusted for false discovery rate using Benjamini-Hochberg method. Full results of the differential abundance analysis can be found in **Table S4**.

To analyse soil microbial community similarity between geographically distant sites from the same ecosystem, we applied the multi-site incidence-based diversity metric zeta diversity (33, 140). This index quantifies the average number of taxa or richness shared across multiple sites (zeta order ζ). Biological replicates of each site were pooled and rarefied at 400 reads per sample for the analysis. Zeta decline, which compares the average number of shared taxa between two to eight sites (ζ_2_ - ζ_8_) from each ecosystem category, was computed using the function *Zeta.decline.mc* in zetadiv v1.2.1 (141) with 500 bootstraps. Values were normalized by the Jaccard method to account for sample richness differences. A power law regression was applied to visualize zeta decline pattern. To ensure robustness, the analysis was carried out on *rplB* OTUs at the taxonomic resolutions of genus, family and order, and on datasets that were unrarefied and rarefied at 300 and 500 counts. The same analysis was repeated with the marker gene *rplP*. Zeta decay, which compares the decrease of shared richness against mean geographic distance between two to six sites (ζ_2_ – ζ_6_) from each ecosystem category, was computed using the function *Zeta.decay* with 500 bootstraps. Unrarefied and rarefied (at 400 counts) datasets of *rplB* and *rplP* at genus levels were analysed. Phylogeny-and abundance-aware beta diversity of sites from the same ecosystem category was computed using weighted UniFrac distances implemented in SingleM v.0.13.2 and ExpressBetaDiversity v1.0.10 (142).

### Metabolic annotation and genome analysis

To holistically estimate the metabolic capability of the soil communities and their ability to utilise organic and inorganic substrates, metagenomes were searched against the KEGG prokaryotic protein database (accessed 22 November 2021) (40), the Carbohydrate-Active enZYmes (CAZy) in dbCAN2 database v10 (143), the MEROPS peptidase database release 12.1 (144), and a custom protein database of representative metabolic markers (https://github.com/GreeningLab/GreeningLab-database) (21). The custom database consists of 52 marker genes involved in carbon fixation (RbcL, AcsB, AclB, Mcr, HbsT, HbsC), hydrogen cycling (large subunit of [NiFe], [FeFe], [Fe]-hydrogenases), carbon monoxide oxidation (CoxL, CooS), methane cycling (McrA, MmoA, PmoA), isoprene oxidation (IsoA), phototrophy (PsaA, PsbA, energy-converting microbial rhodopsin), sulfur cycling (AsrA, FCC, Sqr, DsrA, Sor, SoxB), nitrogen cycling (AmoA, HzsA, NifH, NarG, NapA, NirS, NirK, NrfA, NosZ, NxrA, NorB, Nod), iron cycling (Cyc2, MtrB, OmcB), arsenic cycling (ARO, ArsC), selenium cycling (YgfK), formate oxidation (FdhA), reductive dehalogenation (RdhA), and the respiratory chain (NuoF, SdhA, FrdA, CoxA, CcoN, CyoA, CydA, AtpA). To allow fair comparison of metagenomes from various sources, only quality-filtered forward reads longer than 130 bp and right-trimmed to a maximum 150 bp by SeqKit v2.0.0 (145) were used for the analysis. Metagenomic reads were searched against the gene databases using DIAMOND v.2.0.11 (query cover > 80%) (146). Results were filtered based on an identity threshold of 50%, except for AmoA, NxrA, CoxL, MmoA, [FeFe] and group 4 [NiFe]-hydrogenases, RbcL, and NuoF (all 60%), AtpA, ARO, IsoA, PsbA, and YgfK (all 70%), HbsT (75%), and PsaA (80%) (21). Subgroup and family classification of reads was based on the closest match to the sequences in databases. Read counts for each gene were normalized to reads per kilobase per million (RPKM) by dividing the read count by the total number of reads (in millions) and gene length (in kilobases). Reads were also screened for the 14 universal microbial single copy ribosomal marker genes used in SingleM and PhyloSift (147) by DIAMOND (query cover > 80%, bitscore > 40) and normalized as above. The average gene copy number of a gene in the community was then estimated by dividing RPKM value of the gene by the geomean of RPKM value of the 14 universal single copy ribosomal marker genes. We applied the same method to calculate community 16S rRNA gene copy numbers for normalizing 16S rRNA gene qRCR measurement to cell count. RPKM values for 16S rRNA genes were estimated by 1) mapping quality-filtered reads against the masked and curate SSU NR96 database from phyloFlash v3.4 using MMSeqs2 release 13-45111 easy-search (nucleotide search, query cover > 80%, identity > 70%, sensitivity: 6) (148); 2) removing hits to eukaryotes, mitochondria and chloroplasts; and 3) dividing against an average gene length of 16S rRNA gene (1500 bp) and total number of reads. The applied coverage and identity cutoffs followed the benchmarking results of phyloFlash with regards to the optimal balance in false discovery and sensitivity (133). Statistical difference of the KEGG Orthology (KO) abundance between desert ecosystems and non-desert environments was computed using Welch’s T test with *p*-values adjusted by Benjamini-Hochberg method (**Table S7**).

DRAM v1.2.4 (149) was used to perform metabolic annotations of MAGs, with dbCAN2 database, MEROPS peptidase database, and KEGG protein database, in addition to the search of ribosomal and transfer RNA. Open reading frames (ORFs) in MAGs annotated as peptidases were further analysed through PSORTb v3.0.3 (150) to determine protein subcellular localization and identify potential digestive peptidases. CAZy and peptidase gene copy numbers in MAG were normalized against the genome completeness whereas their proportion in genome was calculated by dividing the total nucleotide length against genome size. In addition, all MAGs were also screened for the presence of metabolic marker genes in our custom database. MAG annotation results can be found in **Table S6**.

### Phylogenetic analysis

Phylogenomic trees were constructed to visualize the phylogenetic breadth of MAGs recovered from the global desert datasets. We adopted the method of the GTDB team to align with their taxonomy inference (151). A concatenated multiple protein sequence alignment was built using the 120 GTDB bacterial marker genes in all bacterial MAGs by GTDB-Tk v1.4.0. IQ-TREE v2.2.0 (152) was then used to construct a maximum likelihood phylogenetic tree with 1000 ultrafast bootstrap replicates (153) under the WAG model of protein evolution with gamma-distributed rate heterogeneity (WAG + G20). The tree was rooted between Gracilicutes and Terrabacteria according to Coleman and Davin *et al.* (154).

### Measurement of carbon and nitrogen stable isotope at natural abundance level

Stable isotope measurement was performed to probe the possible origins of soil nitrogen, total carbon, and organic matter. Approximately 1 g of individual soil was dried at 60°C for 24 h and powder homogenized with a clean mortar and pestle. The ground powder sample was further dried at 60°C overnight. To determine stable isotope ratio of total nitrogen (^15^N /^14^N) and carbon (^13^C/^12^C), an aliquot of 100 mg of sample was weighed on a precision balance (± 0.001 mg) and transferred to a 12 mm × 6 mm tin capsule (Sercon Ltd., UK). For measurement of carbon stable isotope ratio in organic fractions, an aliquot of 100 mg of sample was weighed in an 8 mm × 5 mm silver capsule (Sercon Ltd., UK). Carbonate in the sample was removed through repeated addition of 10% HCl and drying at 60°C until no further effervescence was observed. The dried sample was encapsulated within a tin capsule and all encapsulated samples were pelletized and analysed in the Natural Abundance & IRMS Laboratory at Monash University. The δ^13^C and δ^15^N were measured by combusting the sample at 1050°C in an ANCA GSL2 elemental analyser interfaced with a Hydra 20-22 continuous flow isotope ratio mass spectrometer (CF-IRMS; Sercon Ltd. UK). To ensure accuracy of the isotopic values, internal standards (i.e. sucrose, gelatine and bream) were run concurrently with the samples. These internal standards have been calibrated against internationally recognised reference materials (i.e. USGS 40, USGS41, and IAEA C-6). The analysis additionally yielded information of total nitrogen content, carbon content, and organic carbon fraction for each soil. The machine has a limit of quantification of 0.1 mg for carbon and 0.03 mg for nitrogen, a precision of 0.5 µg for both carbon and nitrogen determination, and a precision of ± 0.1‰ for δ^13^C and ± 0.2‰ δ^15^N.

### Trace gas consumption experiment

Soil microcosms coupled with high sensitivity gas chromatography measurement were set up to determine the capacity of soil microbial communities to oxidize trace gases H_2_, CO, and CH_4_ under dry (native sampling hydration condition) and wet condition (rehydration through rain simulation). For each sample in biological triplicate, 2 g of soil was transferred to a sterile 120-ml serum vial and incubated at either 10°C (for polar deserts and Drass Valley samples) or 20°C (for the rest) in the dark. For the wet condition, 50% v/w of sterile water was added. Samples were allowed to equilibrate with the experimental conditions for 12 h before crimp-sealing the vials with butyl rubber stoppers. Butyl rubber stoppers throughout all trace gas experiments were pre-treated with 0.1 N hot NaOH solution according to the description by Nauer et al. (155) to reduce abiotic emissions of H_2_ and CO from the stopper. An initial mixing ratio of approximately 10 parts per million (ppmv) each of H_2_, CO, and CH_4_ were prepared (via a mixed gas cylinder containing 0.1 % v/v H_2_, CO, and CH_4_ each in N_2_, BOC Australia) and 15 ml of air was applied in the ambient air headspace to reduce the development of under-pressure during repeated sampling. Headspace H_2_, CO, and CH_4_ concentrations were monitored using a VICI gas chromatographic machine with a pulsed discharge helium ionization detector (model TGA-6791-W-4U-2, Valco Instruments Company Inc.) and an autosampler as previously described (156). The machine was regularly calibrated against ultra-pure H_2_, CO, and CH_4_ standards down to the limit of quantification (H_2_: 20 ppbv; CO: 9 ppbv; CH_4_: 500 ppbv). For each condition, heat-killed soil of each biological triplicate (treated at 121°C, 15 p.s.i. for 60 mins) and empty vials (technical duplicate) were included as negative controls.

For calculation of kinetic parameters, time points with individual gas concentrations over 0.4 ppmv were used. First order reaction rate constants were calculated by fitting an exponential model as determined by the lowest overall Akaike information criterion value when compared to a linear model. Actual reaction rate constants of the sample were obtained by correcting against means of negative controls to account for deviations caused by repeated samplings and only resultant values higher than the magnitude of measurement errors of negative controls were retained. Soil atmospheric gas oxidation rates were calculated with respect to the mean atmospheric mixing ratio of the corresponding trace gases (H_2_: 0.53 ppmv; CO: 0.09 ppmv; CH_4_: 1.9 ppmv) (157–159). Assuming all cells are active, cell-specific gas oxidation rates were then inferred by dividing estimated soil cell abundance and the proportion of corresponding gas oxidizers from metagenomic data.

### Respiration assay

Respiration rates of soil communities under dry and wet conditions were measured through an *ex situ* microcosm experiment. Microcosms for each sample in biological triplicates were prepared in sterile 120-ml serum vials with identical incubation temperatures, hydration treatment, and pre-incubation conditions as described for the trace gas consumption experiment, except that 4 g of soil was used. Negative controls using empty vials were prepared. After sealing of the vial, the headspace was flushed with pressurized air (Industrial grade, BOC Australia) at 1 bar for 2 min to achieve a stable initial level of CO_2_ and O_2_. Headspace CO_2_ and N_2_O production was monitored up to 8 days using a VICI gas chromatographic machine as described previously. Daily emission of CO_2_ was quantified by linear regression and corrected against the empty vial controls.

### 14C radioisotope labelling assay

A tracer experiment using radiolabelled carbon dioxide (^14^CO_2_ / H^14^CO_3_ ^-^) was carried out to determine soil chemosynthetic and photosynthetic carbon fixation capacities, given this allows a sensitive detection of carbon incorporation without excessive incubation duration. The biological triplicate was pooled for each location and 0.25 g of soil was transferred to a sterile 5-ml glass vial (MicroAnalytix). The sample was added with 50% v/w of sterile water containing 0.0833 µmol of NaH^14^CO_3_ (equivalent amount as 400 ppmv headspace ^14^CO_2_; Perkin Elmer, 56.6 mCi mmol^-1^) and the vial was immediately sealed with PTFE/silicone septum lid (Supelco, Sigma-Aldrich). The samples were then incubated for 72 h at either 10°C (for polar deserts and Drass Valley samples) or 20°C (for the rest) under three conditions: 1) dark (covered in aluminium foil), 2) light (40 μmol photons m^-2^ s^-1^ under constant illumination), and 3) dark hydrogenotrophic condition (100 ppmv of headspace H_2_). Samples from the Atacama Desert were incubated for 148 h to obtain enough signal. For each condition and location, a technical triplicate and a heat-killed soil control was prepared. Technical duplicate of sample-free incubation control was also included for each condition to monitor background signal. Following incubation, each sample was treated with 2 ml of 1M HCl to remove unfixed CO_2_ and the content was transferred to a 20-ml scintillation vial. After overnight acidification treatment, the sample was added with an additional 1 ml of 1M HCl and allowed to dry at 60°C under a heat lamp. 20 ml of liquid scintillation cocktail (EcoLume™, MP Biomedical) was added to the dried sample and the signal of fixed ^14^C was measured on an automated liquid scintillation counter (Tri-Carb 2810 TR, Perkin Elmer) for 5 min. The machine was regularly calibrated with ^14^C standards of known activity. The amount of ^14^C fixed in each sample was calculated by subtracting scintillation counts from heat-killed controls and converting resultant counts against the reported specific radioactivity of the original bicarbonate solution.

### Driver analysis

Random forest and Spearman correlation analysis were performed to determine the effects of (1) climatological conditions on microbial taxa and gene abundance across soil ecosystems, (2) climatological conditions on physicochemical parameters measured in desert soils, and (3) edaphic conditions on microbial cell abundance and activities measured in desert soils. For (1), multicollinearity of 31 climatological parameters (**Table S1, S3**) was assessed by Spearman correlation using the base R function *cor*. Highly correlated pairs (|*ρ|* > 0.7) were manually inspected and the most ecological significant parameter based on established knowledge was chosen as the predictor, leading to eight climatological predictors (**Fig. 4**). For example, annual mean temperature was retained among other highly correlated parameters such as solar radiation and max/min temperature of warmest/coldest month; similarly, aridity index was retained from highly correlated precipitation measures (e.g. annual mean precipitation) and leaf vegetation index. KO, CAZy (glycoside hydrolase, polysaccharide lyase, carbohydrate esterase, auxiliary activities), and metabolic marker genes with a non-zero abundance in >90% of all soil metagenomes were included for analysis whereas microbial taxa at the genus level based on *rplB* with a minimum occupancy of 50 were used. The importance of climatological parameters in predicting abundance of each gene was determined by random forest analysis using the *rfPermute* function (ntree = 1000, num.rep = 100, na.action = na.omit) in R package rfPermute v2.5.5. Relative importance of each predictor was normalized by dividing %IncMSE (percentage increase in mean squared error) for each predictor by the sum of %IncMSE. Spearman correlation between climatological predictors and abundance of each gene was performed using the base R function *cor* (use = “pairwise.complete.obs). All *p* values were adjusted for false discovery rate using Benjamini-Hochberg method with base R function *p.adjust*. For (2), 29 physicochemical parameters measured in desert soils were included for the analysis (**Table S1**) using the same method. For (3), 15 representative physicochemical predictors were chosen following the above approach to minimize multicollinearity between variables (|*ρ|* < 0.7) and their effects on 12 microbial activity parameters (cell abundance, respiration, atmospheric H_2_, CO, and CH_4_ oxidation at native and wetted hydration conditions, and dark, hydrogenotrophic, and photosynthetic carbon fixation) were determined using both random forest analysis and Spearman correlation analysis. An additional predictor based on cell abundance of microbial mediator of the activity was included to assess relative importance of biological drivers, namely all cells for respiration, H_2_ oxidizers for H_2_ oxidation, CO oxidizers for CO oxidation, CH_4_ oxidizers for CH_4_ oxidation, all autotrophs for dark carbon fixation, CBB-autotrophs for hydrogenotrophic carbon fixation, and photoautotrophs for photosynthetic carbon fixation. Abundance of trace gas oxidizers and autotroph was estimated by multiplying cell count and average gene copy numbers divided by 100 (assuming marker genes are single copy). Random forest and Spearman correlation results are summarized in **Table S9**.

## Data Availability

All metagenomes, metagenomic assemblies, and metagenome-assembled genomes of the desert samples were deposited to the National Center for Biotechnology Information Sequence Read Archive under the BioProject PRJNA832417. Metagenome-assembled genomes reconstructed from the public soil data were deposited to Monash Bridges (doi: 10.26180/24302833). All other study data are included in the article and/or supporting information.

## Acknowledgements

This study was supported by the Australian Research Council (DE170100310 to C.G.; DE250101210 to P.M.L.; DE230101346 to S.K.B.; SR200100005 to S.L.C and C.G.), Australian Antarctic Division (4592 to C.G. and S.L.C.), South African National Antarctic Program (110730 to D.A.C.), National Health & Medical Research Council (APP1178715 salary for C.G.), Antarctic Science Foundation (Traversing the COVID Gap grant 2022 to P.M.L.), Academy of Finland (287545 & 350672 to H.J. & K.M.-M.), National Agency for Research and Development of Chile (Fondecyt project N°1231507 to C.D. and C.G.), Danish National Research Foundation (Center for Permafrost: CENPERM DNRF100 to B.E.; Center for Volatile Interactions: VOLT DNRF168 to A.P.), Spanish National Research Agency (PID2023-147027NB-I00B to A.d.R), Foundational Biodiversity Information Program (FBIP; FBIS160422162807 to J.-B.R.), and Singapore Ministry of Education and Yale-NUS College (R-607-265-331-121 to S.D.J.A.). We thank the academic research discount offered by the EAL, a Southern Cross University National Association of Testing Authorities ISO17025 accredited commercial and research support facility. We greatly appreciate the MonARCH and MASSIVE M3 HPC facility for providing computation platforms and resources, Bethany Coulthard, Vanessa Buzzard and Katie Robins for field and logistic assistance, Steve Pointing for contributing to sample provision, Ros Gleadow and John Beardall for allowing access to the liquid scintillation spectrometer, Rachael Lappan, Guy Shelley and Philipp Nauer for script writing support, Alicia Quinn, Ning Hall and Vera Eate for technical assistance, and Eleonora Chiri, Rhys Grinter, Ya-Jou Chen, Belinda Ferrari and Angelique Ray for helpful discussions.

## Author contributions

C.G. and P.M.L. conceived and supervised the study. P.M.L., C.G., and S.K.B. designed experiments. P.M.L. and C.G. analysed data and wrote the manuscript with inputs from all authors. Different authors were responsible for field sampling and logistical support (S.D.J.A., D.A.C., C.G., S.K.B., A.C., J-B.R., L.K.M., G.Y.A.T., N.W., B.E., A.P., J.B., A.d.R., P.G., H.J., M-M.K, O.G., D.W.G., S.D.S., I.D.H, K.A.W-R., J.D., S.L.C., T.J., P.M.L., T.M., B.F., C.D.), DNA extraction and quantitative PCR assay (P.M.L., T.J.), metagenomic sequencing, assembly and binning (P.M.L.), genome analysis (P.M.L.), metabolic annotation (P.M.L., C.G.), community analysis (P.M.L., S.K.B.), phylogenetic analysis (P.M.L.), geographical and physicochemical analysis (P.M.L.), driver analysis (P.M.L., M.D., C.G.), stable isotope measurement (W.W.W., P.M.L.), trace gas microcosm experiment (P.M.L.), respiration assays (P.M.L., W.W.W.), and ^14^C radioisotope assay (P.M.L., S.K.B., T.N.).

## References

1. M. Cherlet, et al., Eds., World atlas of desertification, 3rd Ed. (Publication Office of the European Union, 2018).

2. D. A. Keith, et al., A function-based typology for Earth’s ecosystems. Nature 610, 513–518 (2022).

3. T. P. Makhalanyane, et al., Microbial ecology of hot desert edaphic systems. FEMS Microbiology Reviews 39, 203–221 (2015).

4. P. M. Leung, et al., Energetic basis of microbial growth and persistence in desert ecosystems. mSystems 5, e00495–19 (2020).

5. F. T. Maestre, et al., Increasing aridity reduces soil microbial diversity and abundance in global drylands. Proceedings of the National Academy of Sciences of the United States of America 112, 15684–15689 (2015).

6. J. W. Neilson, et al., Significant impacts of increasing aridity on the arid soil microbiome. mSystems 2, e00195–16 (2017).

7. D. Schulze-Makuch, et al., Transitory microbial habitat in the hyperarid Atacama Desert. Proceedings of the National Academy of Sciences of the United States of America 115, 2670–2675 (2018).

8. S. K. Bay, et al., Chemosynthetic and photosynthetic bacteria contribute differentially to primary production across a steep desert aridity gradient. The ISME Journal 15, 3339–3356 (2021).

9. A. Rego, et al., Actinobacteria and Cyanobacteria diversity in terrestrial Antarctic microenvironments evaluated by culture-dependent and independent methods. Frontiers in Microbiology 10, 1018 (2019).

10. K. A. Warren-Rhodes, et al., Subsurface microbial habitats in an extreme desert Mars-analog environment. Frontiers in Microbiology 10, 69 (2019).

11. M. Potts, Desiccation tolerance of prokaryotes. Microbiological Reviews 58, 755–805 (1994).

12. J. M. Wood, Bacterial responses to osmotic challenges. Journal of General Physiology 145, 381–388 (2015).

13. T. Ophir, D. L. Gutnick, A role for exopolysaccharides in the protection of microorganisms from desiccation. Applied and Environmental Microbiology 60, 740–745 (1994).

14. Y. Tamaru, Y. Takani, T. Yoshida, T. Sakamoto, Crucial role of extracellular polysaccharides in desiccation and freezing tolerance in the terrestrial cyanobacterium *Nostoc commune*. Applied and Environmental Microbiology 71, 7327–7333 (2005).

15. M. van de Mortel, L. J. Halverson, Cell envelope components contributing to biofilm growth and survival of *Pseudomonas putida* in low-water-content habitats. Molecular Microbiology 52, 735–750 (2004).

16. T. Romantsov, Z. Guan, J. M. Wood, Cardiolipin and the osmotic stress responses of bacteria. Biochimica et Biophysica Acta (BBA) - Biomembranes 1788, 2092–2100 (2009).

17. M. M. Cox, J. R. Battista, *Deinococcus radiodurans* — the consummate survivor. Nature Reviews Microbiology 3, 882–892 (2005).

18. Q. Gao, F. Garcia-Pichel, Microbial ultraviolet sunscreens. Nature Reviews Microbiology 9, 791–802 (2011).

19. P. H. Lebre, P. D. Maayer, D. A. Cowan, Xerotolerant bacteria: surviving through a dry spell. Nature Reviews Microbiology 15, 285–296 (2017).

20. M. Ji, et al., Atmospheric trace gases support primary production in Antarctic desert surface soil. Nature 552, 400–403 (2017).

21. M. Ortiz, et al., Multiple energy sources and metabolic strategies sustain microbial diversity in Antarctic desert soils. Proceedings of the National Academy of Sciences of the United States of America 118, e2025322118 (2021).

22. N. B. Dragone, et al., Elevational constraints on the composition and genomic attributes of microbial communities in Antarctic soils. mSystems 7, e0133021 (2022).

23. A. E. Ray, et al., Atmospheric chemosynthesis is phylogenetically and geographically widespread and contributes significantly to carbon fixation throughout cold deserts. The ISME Journal 16, 2547–2560 (2022).

24. K. Jordaan, et al., Hydrogen-oxidizing bacteria are abundant in desert soils and strongly stimulated by hydration. mSystems 5, e01131–20 (2020).

25. S. Li, et al., Reduced trace gas oxidizers as a response to organic carbon availability linked to oligotrophs in desert fertile islands. The ISME Journal 17, 1257–1266 (2023).

26. S. Imminger, et al., Survival and rapid resuscitation permit limited productivity in desert microbial communities. Nature Communications 15, 3056 (2024).

27. D. V. Meier, S. Imminger, O. Gillor, D. Woebken, Distribution of mixotrophy and desiccation survival mechanisms across microbial genomes in an arid biological soil crust community. mSystems 6, e00786–20 (2021).

28. Y. Hwang, et al., Leave no stone unturned: individually adapted xerotolerant Thaumarchaeota sheltered below the boulders of the Atacama Desert hyperarid core. Microbiome 9, 234 (2021).

29. S. Nayfach, et al., A genomic catalog of Earth’s microbiomes. Nature Biotechnology 39, 499–509 (2021).

30. M. Bahram, et al., Structure and function of the global topsoil microbiome. Nature 560, 233–237 (2018).

31. L. P. Coelho, et al., Towards the biogeography of prokaryotic genes. Nature 601, 252–256 (2022).

32. J. Huang, H. Yu, X. Guan, G. Wang, R. Guo, Accelerated dryland expansion under climate change. Nature Climate Change 6, 166–171 (2016).

33. C. Hui, M. A. McGeoch, Zeta diversity as a concept and metric that unifies incidence-based biodiversity patterns. American Naturalist 184, 684–694 (2014).

34. M. A. McGeoch, et al., Measuring continuous compositional change using decline and decay in zeta diversity. Ecology 100 (2019).

35. N. Fierer, et al., Cross-biome metagenomic analyses of soil microbial communities and their functional attributes. Proceedings of the National Academy of Sciences of the United States of America 109, 21390–21395 (2012).

36. H. Liu, et al., Warmer and drier ecosystems select for smaller bacterial genomes in global soils. iMeta 2, e70 (2023).

37. S. J. Giovannoni, J. Cameron Thrash, B. Temperton, Implications of streamlining theory for microbial ecology. ISME Journal 8, 1553–1565 (2014).

38. A. Rodríguez-Gijón, et al., A genomic perspective across Earth’s microbiomes reveals that genome size in Archaea and Bacteria is linked to ecosystem type and trophic strategy. Frontiers in Microbiology 12, 761869 (2022).

39. T. E. Brewer, K. M. Handley, P. Carini, J. A. Gilbert, N. Fierer, Genome reduction in an abundant and ubiquitous soil bacterium “*Candidatus* Udaeobacter copiosus.” Nature Microbiology 2, 16198 (2016).

40. M. Kanehisa, S. Goto, KEGG: Kyoto Encyclopedia of Genes and Genomes. Nucleic Acids Research 28, 27–30 (2000).

41. P. Siguier, E. Gourbeyre, M. Chandler, Bacterial insertion sequences: their genomic impact and diversity. FEMS Microbiology Reviews 38, 865–891 (2014).

42. A. Ullastres, M. Merenciano, L. Guio, J. González, “Transposable elements: a toolkit for stress and environmental adaptation in bacteria” in Stress and Environmental Regulation of Gene Expression and Adaptation in Bacteria, (John Wiley & Sons, Ltd, 2016), pp. 137–145.

43. A. J. Weisberg, J. H. Chang, Mobile genetic element flexibility as an underlying principle to bacterial evolution. Annu. Rev. Microbiol. 77, 603–624 (2023).

44. K. E. Williamson, J. J. Fuhrmann, K. E. Wommack, M. Radosevich, Viruses in soil ecosystems: an unknown quantity within an unexplored territory. Annu. Rev. Virol. 4, 201–219 (2017).

45. T. P. Makhalanyane, et al., Microbial ecology of hot desert edaphic systems. FEMS Microbiology Reviews 39, 203–221 (2015).

46. Bentley R, Meganathan R, Biosynthesis of vitamin K (menaquinone) in bacteria. Microbiological Reviews 46, 241–280 (1982).

47. M. Boersch, S. Rudrawar, G. Grant, M. Zunk, Menaquinone biosynthesis inhibition: a review of advancements toward a new antibiotic mechanism. RSC Adv 8, 5099–5105 (2018).

48. R. Duman, et al., Structural and genetic analyses reveal the protein SepF as a new membrane anchor for the Z ring. Proceedings of the National Academy of Sciences 110, E4601–E4610 (2013).

49. E. Sauvage, F. Kerff, M. Terrak, J. A. Ayala, P. Charlier, The penicillin-binding proteins: structure and role in peptidoglycan biosynthesis. FEMS Microbiology Reviews 32, 234–258 (2008).

50. N. Pishesha, J. R. Ingram, H. L. Ploegh, Sortase A: a model for transpeptidation and its biological applications. Annu. Rev. Cell Dev. Biol. 34, 163–188 (2018).

51. M. S. Prasad, R. P. Bhole, P. B. Khedekar, R. V. Chikhale, *Mycobacterium* enoyl acyl carrier protein reductase (InhA): A key target for antitubercular drug discovery. Bioorganic Chemistry 115, 105242 (2021).

52. C. Coleine, et al., Dryland microbiomes reveal community adaptations to desertification and climate change. The ISME Journal 18, wrae056 (2024).

53. H. Shaomei, et al., Patterns in wetland microbial community composition and functional gene repertoire associated with methane emissions. mBio 6, e00066–15 (2015).

54. T. Wagner, C. R. Magill, J. O. Herrle, “Carbon isotopes” in Encyclopedia of Geochemistry: A Comprehensive Reference Source on the Chemistry of the Earth, W. M. White, Ed. (Springer International Publishing, 2018), pp. 194–204.

55. P. J. Thomas, et al., Isotope discrimination by form IC RubisCO from *Ralstonia eutropha* and *Rhodobacter sphaeroides*, metabolically versatile members of ‘Proteobacteria’ from aquatic and soil habitats. Environmental Microbiology 21, 72–80 (2019).

56. C. R. Edwards, et al., Draft genome sequence of uncultured upland soil cluster Gammaproteobacteria gives molecular insights into high-affinity methanotrophy. Genome Announcements 5, e00047–17 (2017).

57. H. Koch, et al., Growth of nitrite-oxidizing bacteria by aerobic hydrogen oxidation. Science 345, 1052–1054 (2014).

58. P. M. Leung, et al., A nitrite-oxidising bacterium constitutively consumes atmospheric hydrogen. The ISME Journal 16, 2213–2219 (2022).

59. I. Noy-Meir, Desert ecosystems: environment and producers. Annual Review of Ecology and Systematics 4, 25–51 (1973).

60. R. G. Strieg, T. A. McConnaughey, D. C. Thorstenson, E. P. Weeks, J. C. Woodward, Consumption of atmospheric methane by desert soils. Nature 357, 145–147 (1992).

61. J. P. Schimel, Life in dry soils: Effects of drought on soil microbial communities and processes. *Annual Review of Ecology*, Evolution, and Systematics 49, 409–432 (2018).

62. G. M. King, “Water potential as a master variable for atmosphere-soil trace gas exchange in arid and semiarid ecosystems” in The Biology of Arid Soils, B. Steven, Ed. (De Gruyter, Berlin, Boston, 2017), pp. 31–45.

63. B. S. Guida, M. Bose, F. Garcia-Pichel, Carbon fixation from mineral carbonates. Nature Communications 8, 1025 (2017).

64. F. Garcia-Pichel, O. Pringault, Microbiology: Cyanobacteria track water in desert soils. Nature 413, 380–381 (2001).

65. H. Raanan, et al., Simulated soil crust conditions in a chamber system provide new insights on cyanobacterial acclimation to desiccation. Environmental Microbiology 18, 414–426 (2016).

66. S. E. Fick, R. J. Hijmans, WorldClim 2: new 1-km spatial resolution climate surfaces for global land areas. International Journal of Climatology 37, 4302–4315 (2017).

67. R. J. Zomer, J. Xu, A. Trabucco, Version 3 of the global aridity index and potential evapotranspiration database. Scientific Data 9, 409 (2022).

68. X. Xu, P. E. Thornton, W. M. Post, A global analysis of soil microbial biomass carbon, nitrogen and phosphorus in terrestrial ecosystems. Global Ecology and Biogeography 22, 737–749 (2013).

69. J. Mao, B. Yan, Global monthly mean leaf area index climatology, 1981-2015 (version 1). ORNL Distributed Active Archive Center. 10.3334/ORNLDAAC/1653.

70. J. E. Pinzon, et al., Global vegetation greenness (NDVI) from AVHRR GIMMS-3G+, 1981-2022 (version 1). ORNL Distributed Active Archive Center. 10.3334/ORNLDAAC/2187.

71. N. Madani, J. S. Kimball, S. W. Running, Improving global gross primary productivity estimates by computing optimum light use efficiencies using flux tower data. Journal of Geophysical Research: Biogeosciences 122, 2939–2951 (2017).

72. A. E. Parada, D. M. Needham, J. A. Fuhrman, Every base matters: Assessing small subunit rRNA primers for marine microbiomes with mock communities, time series and global field samples. Environmental Microbiology 18, 1403–1414 (2016).

73. A. Apprill, S. Mcnally, R. Parsons, L. Weber, Minor revision to V4 region SSU rRNA 806R gene primer greatly increases detection of SAR11 bacterioplankton. Aquatic Microbial Ecology 75, 129–137 (2015).

74. S. Nurk, D. Meleshko, A. Korobeynikov, P. A. Pevzner, MetaSPAdes: A new versatile metagenomic assembler. Genome Research 27, 824–834 (2017).

75. D. Li, C. M. Liu, R. Luo, K. Sadakane, T. W. Lam, MEGAHIT: An ultra-fast single-node solution for large and complex metagenomics assembly via succinct de Bruijn graph. Bioinformatics 31, 1674–1676 (2015).

76. B. Langmead, S. L. Salzberg, Fast gapped-read alignment with Bowtie 2. Nature Methods 9, 357–359 (2012).

77. I. J. Miller, et al., Autometa: automated extraction of microbial genomes from individual shotgun metagenomes. Nucleic Acids Research 47, e57 (2019).

78. J. Alneberg, et al., Binning metagenomic contigs by coverage and composition. Nature methods 11, 1144–1146 (2014).

79. Y.-W. Wu, B. A. Simmons, S. W. Singer, MaxBin 2.0: an automated binning algorithm to recover genomes from multiple metagenomic datasets. Bioinformatics 32, 605–607 (2015).

80. D. Kang, et al., MetaBAT 2: an adaptive binning algorithm for robust and efficient genome reconstruction from metagenome assemblies. PeerJ 7, e7359 (2019).

81. C. M. K. Sieber, et al., Recovery of genomes from metagenomes via a dereplication, aggregation and scoring strategy. Nature microbiology 3, 1 (2018).

82. D. H. Parks, et al., Recovery of nearly 8,000 metagenome-assembled genomes substantially expands the tree of life. Nature Microbiology 2, 1533–1542 (2017).

83. M. R. Olm, C. T. Brown, B. Brooks, J. F. Banfield, dRep: a tool for fast and accurate genomic comparisons that enables improved genome recovery from metagenomes through de-replication. The ISME journal 11, 2864 (2017).

84. D. H. Parks, M. Imelfort, C. T. Skennerton, P. Hugenholtz, G. W. Tyson, CheckM: Assessing the quality of microbial genomes recovered from isolates, single cells, and metagenomes. Genome Research 25, 1043–1055 (2015).

85. R. M. Bowers, et al., Minimum information about a single amplified genome (MISAG) and a metagenome-assembled genome (MIMAG) of bacteria and archaea. Nature Biotechnology 35, 725–731 (2017).

86. P.-A. Chaumeil, A. J. Mussig, P. Hugenholtz, D. H. Parks, GTDB-Tk: a toolkit to classify genomes with the Genome Taxonomy Database. Bioinformatics 36, 1925–1927 (2020).

87. D. H. Parks, et al., A complete domain-to-species taxonomy for Bacteria and Archaea. Nature Biotechnology 38, 1079–1086 (2020).

88. B. Ma, et al., Genetic correlation network prediction of forest soil microbial functional organization. The ISME Journal 12, 2492–2505 (2018).

89. Y. Zhong, W. Yan, R. Wang, W. Wang, Z. Shangguan, Decreased occurrence of carbon cycle functions in microbial communities along with long-term secondary succession. Soil Biology and Biochemistry 123, 207–217 (2018).

90. R. Starke, et al., Niche differentiation of bacteria and fungi in carbon and nitrogen cycling of different habitats in a temperate coniferous forest: A metaproteomic approach. Soil Biology and Biochemistry 155, 108170 (2021).

91. R. C. Wilhelm, et al., A metagenomic survey of forest soil microbial communities more than a decade after timber harvesting. Scientific Data 4, 170092 (2017).

92. M. B. N. Albright, et al., Short-term transcriptional response of microbial communities to nitrogen fertilization in a pine forest soil. Applied and Environmental Microbiology 84, e00598–18 (2018).

93. C. N. Kelly, G. W. Schwaner, J. R. Cumming, T. P. Driscoll, Metagenomic reconstruction of nitrogen and carbon cycling pathways in forest soil: Influence of different hardwood tree species. Soil Biology and Biochemistry 156, 108226 (2021).

94. F. Schulz, et al., Hidden diversity of soil giant viruses. Nature Communications 9, 4881 (2018).

95. N. C. Dove, M. S. Torn, S. C. Hart, N. Taş, Metabolic capabilities mute positive response to direct and indirect impacts of warming throughout the soil profile. Nature Communications 12, 2089 (2021).

96. P. B. Matheus Carnevali, et al., Meanders as a scaling motif for understanding of floodplain soil microbiome and biogeochemical potential at the watershed scale. Microbiome 9, 121 (2021).

97. M. E. Kroeger, et al., New biological insights into how deforestation in Amazonia affects soil microbial communities using metagenomics and metagenome-assembled genomes. Frontiers in Microbiology 9, 1635 (2018).

98. L. W. Mendes, J. M. Raaijmakers, M. de Hollander, R. Mendes, S. M. Tsai, Influence of resistance breeding in common bean on rhizosphere microbiome composition and function. The ISME Journal 12, 212–224 (2018).

99. Q. Yao, et al., Community proteogenomics reveals the systemic impact of phosphorus availability on microbial functions in tropical soil. Nature Ecology & Evolution 2, 499–509 (2018).

100. Z. Zheng, et al., The composition of antibiotic resistance genes is not affected by grazing but is determined by microorganisms in grassland soils. Science of The Total Environment 761, 143205 (2021).

101. A. A. D. Broadbent, et al., Climate change alters temporal dynamics of alpine soil microbial functioning and biogeochemical cycling via earlier snowmelt. The ISME Journal 15, 2264–2275 (2021).

102. R. Mackelprang, et al., Microbial community structure and functional potential in cultivated and native tallgrass prairie soils of the midwestern United States. Frontiers in Microbiology 9, 1775 (2018).

103. R. C. Taniya, et al., Metaphenomic responses of a native prairie soil microbiome to moisture perturbations. mSystems 4, e00061–19 (2022).

104. E. R. Johnston, et al., Metagenomics reveals pervasive bacterial populations and reduced community diversity across the Alaska tundra ecosystem. Frontiers in Microbiology 7, 579 (2016).

105. A. Crits-Christoph, S. Diamond, C. N. Butterfield, B. C. Thomas, J. F. Banfield, Novel soil bacteria possess diverse genes for secondary metabolite biosynthesis. Nature 558, 440–444 (2018).

106. Y. Julian, et al., Comparative metagenomics reveals enhanced nutrient cycling potential after 2 years of biochar amendment in a tropical oxisol. Applied and Environmental Microbiology 85, e02957–18 (2019).

107. A. P. Camargo, et al., Microbiomes of Velloziaceae from phosphorus-impoverished soils of the campos rupestres, a biodiversity hotspot. Scientific Data 6, 140 (2019).

108. X. Rui, et al., Metabolic potentials of members of the class Acidobacteriia in metal-contaminated soils revealed by metagenomic analysis. Environmental Microbiology 24, 803–818 (2022).

109. B. O. Oluranti, O. O. Pelumi, I. N. Ozede, S. F. J, Survey of maize rhizosphere microbiome using shotgun metagenomics. Microbiology Resource Announcements 10, e01309–20 (2022).

110. S. R. Pratap, J. A. Kumar, D. Meenakshi, S. F. J, Metagenomic analysis of microbial diversity in cotton rhizosphere soil in Alwar, India. Microbiology Resource Announcements 9, e00987–20 (2022).

111. A. Usyskin-Tonne, Y. Hadar, U. Yermiyahu, D. Minz, Elevated CO_2_ and nitrate levels increase wheat root-associated bacterial abundance and impact rhizosphere microbial community composition and function. The ISME Journal 15, 1073–1084 (2021).

112. J. Nelkner, et al., Effect of long-term farming practices on agricultural soil microbiome members represented by metagenomically assembled genomes (MAGs) and their predicted plant-beneficial genes. Genes 10, 424 (2019).

113. V. M. Flores-Núñez, et al., Functional signatures of the epiphytic prokaryotic microbiome of agaves and cacti. Frontiers in Microbiology 10, 3044 (2020).

114. P. F. Chuckran, et al., Metagenomes and metatranscriptomes of a glucose-amended agricultural soil. Microbiology Resource Announcements 9, e00895–20 (2020).

115. C. Santos-Medellin, et al., Viromes outperform total metagenomes in revealing the spatiotemporal patterns of agricultural soil viral communities. The ISME Journal 15, 1956–1970 (2021).

116. L. Ramírez-Fernández, L. H. Orellana, E. R. Johnston, K. T. Konstantinidis, J. Orlando, Diversity of microbial communities and genes involved in nitrous oxide emissions in Antarctic soils impacted by marine animals as revealed by metagenomics and 100 metagenome-assembled genomes. Science of The Total Environment 788, 147693 (2021).

117. V. B. Centurion, et al., Dynamics of microbial stress responses driven by abiotic changes along a temporal gradient in Deception Island, Maritime Antarctica. Science of The Total Environment 758, 143671 (2021).

118. S. Nie, et al., Desulfobacterales stimulates nitrate reduction in the mangrove ecosystem of a subtropical gulf. Science of The Total Environment 769, 144562 (2021).

119. H.-Y. Li, et al., The chemodiversity of paddy soil dissolved organic matter correlates with microbial community at continental scales. Microbiome 6, 187 (2018).

120. Y. Zhong, et al., Soil microbial mechanisms promoting ultrahigh rice yield. Soil Biology and Biochemistry 143, 107741 (2020).

121. B. J. Woodcroft, et al., Genome-centric view of carbon processing in thawing permafrost. Nature 560, 49–54 (2018).

122. A. L. Peralta, et al., Metagenomes from experimental hydrologic manipulation of restored coastal plain wetland soils (Tyrell County, North Carolina). Microbiology Resource Announcements 9, e00882–20 (2020).

123. C. P. Fernández-Baca, et al., Changes in rhizosphere soil microbial communities across plant developmental stages of high and low methane emitting rice genotypes. Soil Biology and Biochemistry 156, 108233 (2021).

124. R. M. Wilson, et al., Soil metabolome response to whole-ecosystem warming at the spruce and peatland responses under changing environments experiment. Proceedings of the National Academy of Sciences of the United States of America 118, e2004192118 (2021).

125. M. Espenberg, et al., Differences in microbial community structure and nitrogen cycling in natural and drained tropical peatland soils. Scientific Reports 8, 4742 (2018).

126. J.-Y. Li, et al., Comparative metagenomics of two distinct biological soil crusts in the Tengger Desert, China. Soil Biology and Biochemistry 140, 107637 (2020).

127. T. L. Swenson, U. Karaoz, J. M. Swenson, B. P. Bowen, T. R. Northen, Linking soil biology and chemistry in biological soil crust using isolate exometabolomics. Nature Communications 9, 1–10 (2018).

128. E. Ertekin, V. Meslier, A. Browning, J. Treadgold, J. DiRuggiero, Rock structure drives the taxonomic and functional diversity of endolithic microbial communities in extreme environments. Environmental Microbiology 23, 3937–3956 (2021).

129. G. Uritskiy, et al., Halophilic microbial community compositional shift after a rare rainfall in the Atacama Desert. The ISME Journal 13, 2737–2749 (2019).

130. U. F. Lingappa, et al., An ecophysiological explanation for manganese enrichment in rock varnish. Proceedings of the National Academy of Sciences of the United States of America 118, e2025188118 (2021).

131. B. J. Woodcroft, et al., Comprehensive taxonomic identification of microbial species in metagenomic data using SingleM and Sandpiper. Nature Biotechnology (2025). 10.1038/s41587-025-02738-1.

132. Y. Lan, G. Rosen, R. Hershberg, Marker genes that are less conserved in their sequences are useful for predicting genome-wide similarity levels between closely related prokaryotic strains. Microbiome 4 (2016).

133. H. R. Gruber-Vodicka, B. K. B. Seah, E. Pruesse, phyloFlash: rapid small-subunit rRNA profiling and targeted assembly from metagenomes. mSystems 5, e00920–20 (2020).

134. C. Quast, et al., The SILVA ribosomal RNA gene database project: improved data processing and web-based tools. Nucleic Acids Research 41, D590–D596 (2013).

135. A. D. Fernandes, et al., Unifying the analysis of high-throughput sequencing datasets: characterizing RNA-seq, 16S rRNA gene sequencing and selective growth experiments by compositional data analysis. Microbiome 2, 15 (2014).

136. H. Lin, S. D. Peddada, Analysis of compositions of microbiomes with bias correction. Nature Communications 11, 3514 (2020).

137. H. Zhou, K. He, J. Chen, X. Zhang, LinDA: linear models for differential abundance analysis of microbiome compositional data. Genome Biology 23, 95 (2022).

138. H. Mallick, et al., Multivariable association discovery in population-scale meta-omics studies. PLOS Computational Biology 17, e1009442 (2021).

139. J. T. Nearing, et al., Microbiome differential abundance methods produce different results across 38 datasets. Nature Communications 13, 342 (2022).

140. S. K. Bay, et al., Soil bacterial communities exhibit strong biogeographic patterns at fine taxonomic resolution. mSystems 5, e00540–20 (2020).

141. G. Latombe, M. A. McGeoch, D. A. Nipperess, C. Hui, zetadiv: An R package for computing compositional change across multiple sites, assemblages or cases. bioRxiv (2018). 10.1101/324897.

142. D. H. Parks, R. G. Beiko, Measures of phylogenetic differentiation provide robust and complementary insights into microbial communities. ISME Journal 7, 173–183 (2013).

143. H. Zhang, et al., dbCAN2: a meta server for automated carbohydrate-active enzyme annotation. Nucleic Acids Research 46, W95–W101 (2018).

144. N. D. Rawlings, et al., The MEROPS database of proteolytic enzymes, their substrates and inhibitors in 2017 and a comparison with peptidases in the PANTHER database. Nucleic Acids Research 46, D624–D632 (2018).

145. W. Shen, S. Le, Y. Li, F. Hu, SeqKit: a cross-platform and ultrafast toolkit for FASTA/Q file manipulation. PLoS ONE 11, e0163962 (2016).

146. B. Buchfink, C. Xie, D. H. Huson, Fast and sensitive protein alignment using DIAMOND. Nature methods 12, 59 (2014).

147. A. E. Darling, et al., PhyloSift: phylogenetic analysis of genomes and metagenomes. PeerJ 2014, e243 (2014).

148. M. Steinegger, J. Söding, MMseqs2 enables sensitive protein sequence searching for the analysis of massive data sets. Nature Biotechnology 35, 1026–1028 (2017).

149. M. Shaffer, et al., DRAM for distilling microbial metabolism to automate the curation of microbiome function. Nucleic Acids Research 48, 8883–8900 (2020).

150. N. Y. Yu, et al., PSORTb 3.0: improved protein subcellular localization prediction with refined localization subcategories and predictive capabilities for all prokaryotes. Bioinformatics 26, 1608–1615 (2010).

151. D. H. H. Parks, et al., A standardized bacterial taxonomy based on genome phylogeny substantially revises the tree of life. Nature Biotechnology 36, 996 (2018).

152. B. Q. Minh, et al., IQ-TREE 2: new models and efficient methods for phylogenetic inference in the genomic era. Molecular Biology and Evolution 37, 1530–1534 (2020).

153. D. T. Hoang, O. Chernomor, A. von Haeseler, B. Q. Minh, L. S. Vinh, UFBoot2: improving the ultrafast bootstrap approximation. Molecular Biology and Evolution 35, 518–522 (2018).

154. G. A. Coleman, et al., A rooted phylogeny resolves early bacterial evolution. Science 372, eabe0511 (2021).

155. P. A. Nauer, E. Chiri, T. Jirapanjawat, C. Greening, P. L. M. Cook, Technical note: inexpensive modification of exetainers for the reliable storage of trace-level hydrogen and carbon monoxide gas samples. Biogeosciences 18, 729–737 (2021).

156. Z. F. Islam, et al., Two Chloroflexi classes independently evolved the ability to persist on atmospheric hydrogen and carbon monoxide. The ISME Journal 13, 1801–1813 (2019).

157. P. C. C. Novelli, et al., Molecular hydrogen in the troposphere: global distribution and budget. Journal of Geophysical Research Atmospheres 104, 30427–30444 (1999).

158. P. C. Novelli, L. P. Steele, P. P. Tans, Mixing ratios of carbon monoxide in the troposphere. Journal of Geophysical Research 97, 20731–20750 (1992).

159. S. Kirschke, et al., Three decades of global methane sources and sinks. Nature Geoscience 6, 813–823 (2013).

