## Supplementary Information for "Aridity drives global convergence of desert microbiomes and biogeochemical activities"

**Supplementary Note 1: Desert community analysis**.

The profiling of microbial community composition in desert and global soils using classified metagenomic reads of the universal single copy ribosomal protein *rplB* and the small subunit (SSU) of rRNA gene reveals a high consistency (**Table S4-5, Fig. S2B-C**). Actinobacteriota, Proteobacteria, Acidobacteriota, and Chloroflexota are the dominant desert phyla (**Fig. 1B**), together accounting for 84% and 69% of classified *rplB* reads and 78% and 69% of SSU rRNA gene reads in hot and cold deserts and polar lands, respectively. Monoderm Actinobacteriota (Thermoleophilia, Actinomycetia, Acidimicrobiia, Rubrobacteria, and *Ca*. Aridivitia (UBA4738) (1)) and Chloroflexota (Chloroflexia, and Limnocylindria) appear to be overrepresented at the class level, together constituting 65% and 36% of *rplB* reads in hot and cold deserts and polar lands, respectively. To systematically compare the abundance of microbial taxa in deserts versus non-desert ecosystems, we performed differential abundance analysis using four independent methods on *rplB* reads (**Materials and Methods; Table S4**). Here, a taxon is considered differentially abundant (or less abundant) if at least two tools called for significance difference in the same direction (adjust $p < 0.05$). Except for Limnocylindria in agricultural and grassland soils, all seven monoderm classes are differential abundant in desert compared to non-desert ecosystems; these classes are also differentially abundant in polar land compared to forest and wetland (**Fig. 1B**). The most dominant archaeal class Nitrososphaeria in the sampled soils are also significantly more abundant in non-polar desert except when compared with agricultural soils where common fertilization practice likely introduces elevated levels of ammonium for ammonia-oxidizing archaea. In contrast, most diderm classes widespread in non-desert soils showed a contrasting pattern, with Gammaproteobacteria, Terriglobia (Acidobacteriae), Vicinamibacteria, Verrucomicrobiae, and Planctomycetia significantly less abundant in non-polar deserts, despite accounting for one-third of non-desert soil communities (**Fig. 1B**). The two diderm classes Gemmatimonadetes and Blastocatellia (Acidobacteriota) are notable exceptions, being more abundant in both non-polar and polar desert soils when compared with forest and wetlands (**Fig. 1B**). In line with previous observations that photoautotrophs are constrained to specific lithic and biocrust niches in deserts, relative abundance of Cyanobacteria in nonpolar deserts (0.19%) and polar lands (0.05%) are 44 and 152 times less than in biocrusts and rocks, respectively (**Table S4**).

Based on the differential abundance trend of dominant taxa, it appears that geographically distant deserts select for compositionally similar communities. We evaluated this hypothesis using the multisite incidence-based diversity metric zeta diversity (2, 3), which quantifies the average number of taxa (or richness following Jaccard normalization) shared across multiple sites. A zeta decline analysis on *rplB* reads at genus level showed that, across combinations of any two to seven sites in nonpolar deserts, the shared richness among the community is always higher than any other ecosystem (**Fig. 1C, Table S4**). This suggests that nonpolar deserts select for common dominant taxa. However, selection for common communities in polar lands is less apparent and zeta decline was of similar magnitude as grassland and forest soils. To account for geographic distance between sites, we analysed distance decay patterns of pairwise and multisite zeta diversity (zeta decay) (**Fig. S2H**). The results consistently showed nonpolar deserts distinctively share the highest common microbial richness across increasing geographic separation. Interestingly, multisite zeta diversity of polar sites did not decrease with increasing distance compared to most other environments, suggesting some taxa are likely consistently selected in this habitat. We additionally analysed pairwise abundance-based beta diversity (weighted UniFrac) to evaluate compositional similarity between sites. Pairwise dissimilarity between sites of nonpolar deserts is significantly lower than all other ecosystems (all $p < 0.0001$, Welch's t-test) and that of polar lands is also significantly lower than other non-desert ecosystems (all $p < 0.05$, Welch's t-test) except for agricultural land (**Fig. 1C**). Highly concordant patterns of zeta and beta diversity were obtained for the alternative marker gene *rplP*, and unrarefied and rarefied datasets at different taxonomic resolutions (**Fig. S2F-I**). While the data suggest a strong selection of dominant taxa at genus and higher ranks in deserts, we acknowledge that a more extensive and structured global survey is necessary to fully evaluate this pattern, especially with regards to deeper sequencing depths and finer taxonomic resolutions.

### Supplementary Note 2: Energy sources supporting desert microbes

The genetic capacity for autotrophy was strongly correlated with hydrogenotrophy and nitrification, but not with other chemosynthetic or photosynthetic processes. The abundance of *rbcL* is positively correlated with the high-affinity [NiFe]-hydrogenases ($R^2$ = 0.80) (**Fig. 3G**) and both are substantially enriched in desert communities. In common with previous findings in Antarctic soils (4) and Negev Desert (5), this suggests that CBB cycle-based chemosynthetic $CO_2$ fixation is primarily fuelled by atmospheric $H_2$ as an electron donor. Copy numbers of *rbcL* (0.44) and hydrogenase (1.45) are the highest in the hyper-arid Atacama Desert (**Fig. 3D**), the site with the strongest $^{13}C$ depletion in soil organic matter (**Fig. 6A**). Gene copy numbers of *hbs* and *aclB* are linearly correlated with ammonia monooxygenase *amoA* ($R^2$ = 0.86) and nitrite oxidoreductase *nxrA* ($R^2$ = 0.68), respectively (**Fig. 3G**), indicating that desert carbon fixation through the 4-hydroxybutyrate cycle and reductive TCA cycle is likely driven by nitrification.

The presence of metabolic markers in MAGs mirrors the gene-centric short read analysis (**Fig. S6B**). High-affinity [NiFe]-hydrogenases and RuBisCO are encoded by 55 orders and 25 orders, respectively. The tight coupling of the two processes for chemolithoautotrophy (termed aerotrophy (6)) is evidenced by that two-thirds of RuBisCO-encoding MAGs co-encode uptake hydrogenases, including multiple Actinobacteriota classes, Dormibacteria, Limnocylindria, Ktedonobacteria, and Burkholderiales (**Fig. 5D, Table S6**). Nitrososphaerales, accounting for 88% of archaeal *rplB* reads in deserts, exclusively co-encodes *amoA* and *hbs*; whereas Nitrospirales exclusively co-encodes *nxrA* and *aclB* (**Fig. 5D**). Their abundance parallels with that of the marker genes mediating these metabolisms (**Fig. 5C, Fig. 3C**), inferring they are respectively the major ammonia- and nitrite-oxidizing taxa in deserts. A desert Nitrospirales MAG (KAP_bin38) also encodes a group 1h [NiFe]-hydrogenase (**Table S6**), suggesting $H_2$ oxidation may also support some nitrifiers as demonstrated in *Nitrospira moscoviensis* (7, 8). Marker genes mediating other autotrophic pathways were not detected in MAGs. Altogether, diverse hydrogenotrophic bacteria and various nitrifiers emerge as major players in desert soil primary production and their autotrophic capabilities may contribute to their dominance across deserts.

Desert microbes also use other atmospheric substrates, inorganic edaphic compounds, and sunlight to meet energy and carbon needs. 48% of the community, spanning diverse monoderm bacteria and some acidobacterial MAGs, encode carbon monoxide dehydrogenases (*coxL*) for atmospheric CO consumption (**Fig. 3D, Fig. 5D**). This gene is also present in 37.5% of *rbcL*-encoding MAGs (**Table S6**), suggesting carboxydotrophic carbon fixation or mixotrophic growth potential in addition to enhancing survival during starvation (9, 10). Although methane oxidation is a relatively rare trait in deserts (*pmoA*/*mmoA*, 0.70%; encoded by *Methylocella* (*Methylocapsa*/USCα) and Methyloligotrophales (JACCXJ01/USCγ) (6) MAGs), it is enriched in comparison to agricultural and grassland soils (**Fig. 3E**). Methyloligotrophales, inferred as atmospheric $CH_4$ oxidizers (1, 11), has the highest relative abundance in the desert community (**Fig. 5B**), revealing novel high-affinity methanotrophs find a niche in oligotrophic deserts and may contribute to carbon sequestration. Together with the prevalence of hydrogenotrophy (58.5% in desert versus 23.9% in non-desert soils), atmospheric trace gases appear to be particularly important energy sources. A significant proportion of bacteria encode genes for the oxidation of sulfide (*fcc*, 6.5%; *sqr*, 6.1%), thiosulfate (*soxB*, 4.2%), ferrous iron (*cyc2*, 2.7%), and arsenite (*aro*, 0.70%) (**Fig. 3D**) spanning 18 diderm and 9 monoderm orders (**Fig. 5D**). These genes are more prevalent in Arctic and sub-Antarctic sites (**Fig. 3D**), potentially reflecting that the higher moisture within these soils enhances the accessibility of these edaphic substrates to microorganisms (**Fig. S1B**). Moreover, the capacity to harvest light through anoxygenic phototrophy (*psaA* 0.35%; *psbA* 2.1%) is enriched in desert soils compared to other ecosystems (**Fig. 3C**), despite the low abundance of Cyanobacteria (0.1%) (**Fig. S1**). Photosystem II is present in three

Proteobacteria MAGs (Acetobacterales, Steroidobacterales, *Rubrivivax*) (**Fig. 5D**). Extending our previous finding in Antarctic Mackay Glacier soils (1), microbial rhodopsin (*rho*) is the most widespread light-harvesting protein in desert communities. Estimated to be present in 1.8% and 5.2% cells in nonpolar deserts and polar lands respectively, this gene is 2.3- and 6.7-fold more enriched compared to non-desert environments (**Fig. 3C-D**). In dryland biocrust and rock samples, copy numbers for *psaA* and *psbA* are 0.21 and 0.73, respectively, whereas nearly one-third of the community harbours a copy of microbial rhodopsin, revealing an overlooked niche of the metabolism outside aquatic habitats. The gene is widespread in MAGs of the abundant order Rubrobacterales while also present in MAGs from seven other orders (**Fig. 5D**).

### Supplementary Figures

**Figure S1. Characterization of climate, soil physicochemistry, nutrient profile and cellular abundance in the 25 hot, cold, and polar desert sites**. (**A**) Geographic and climate variables based on WorldClim database v2.1 (12) and Global Aridity Index and Potential Evapo-Transpiration (ET0) Database v5 (13). Note that aridity index and precipitation modelling were not accurately modelled for polar regions and were excluded. (**B**) Major soil physicochemical and nutrient profiles. (**C**) Mineral and trace metal content. (**D**) Estimated abundance of microbial cells in soils based on 16S rRNA gene qPCR and metagenomic short read analysis. (**E**) A biplot visualizing the principal component analysis of 26 geographic and climate variables (aridity index, elevation, potential evapotranspiration, standard deviation of potential evapotranspiration, solar radiation, water vapor pressure, wind speed and 19 BioClimate variables) of the 25 sites. The two principal component axes together explain 70% of the site climate variation. (**F**) A biplot visualizing the principal component analysis of 27 representative soil physicochemical variables (pH, electrical conductivity, moisture content, total carbon, total organic carbon, total nitrogen, nitrate nitrogen, ammonium nitrogen, total $\delta^{13}C$, total $\delta^{15}N$, organic $\delta^{13}C$, sulfur, effective cation exchange capacity (ECEC), calcium, magnesium, potassium, exchangeable sodium, aluminium, phosphorus (Colwell), manganese, iron, copper, boron, silicon, gravel, sand, silt/clay content) of soils from the 25 sites. The two principal component axes together explain 43% of the soil variation.

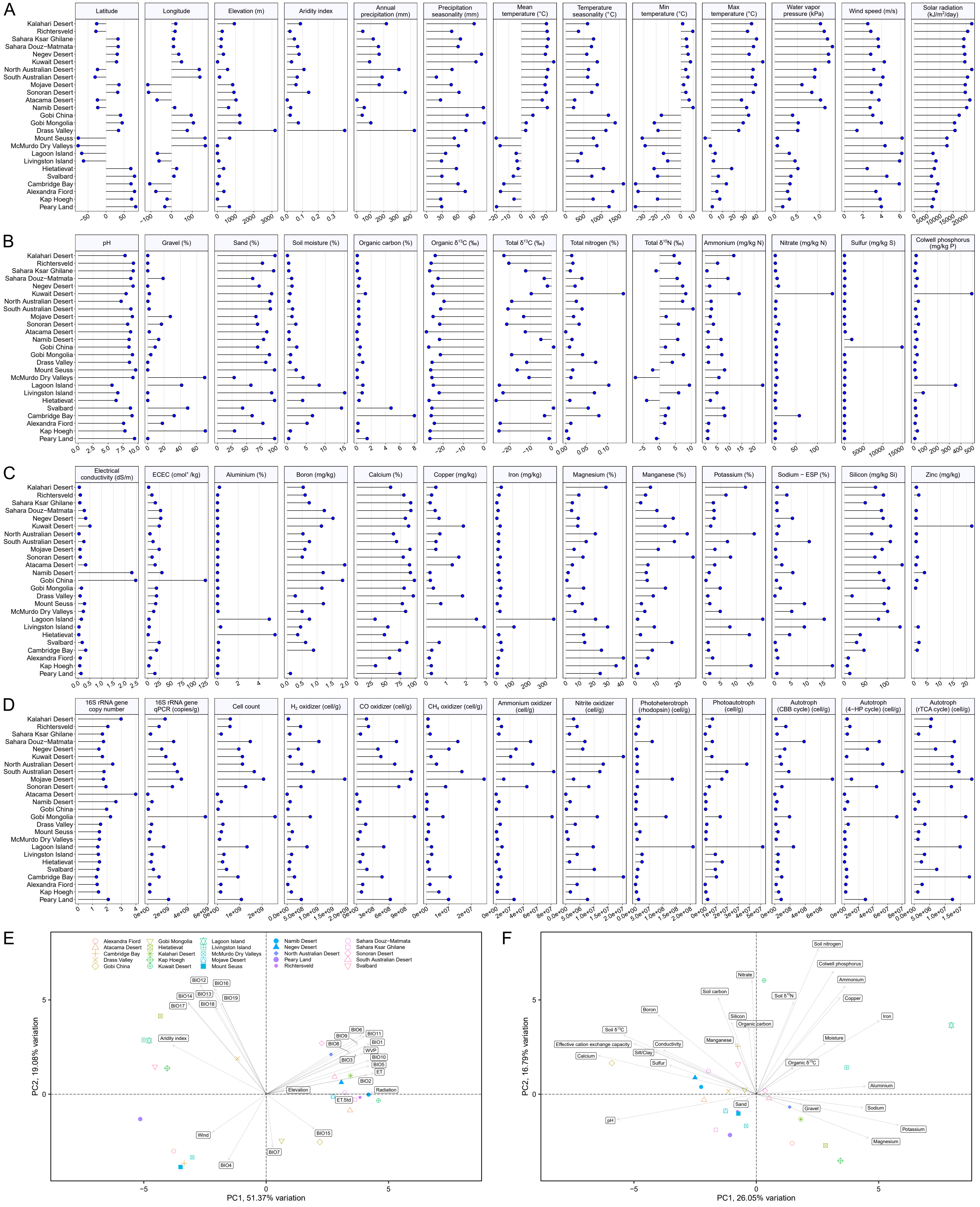

A
Latitude
Longitude
Elevation (m)
Aridity index
Annual precipitation (mm)
Precipitation seasonality (mm)
Mean temperature (°C)
Temperature seasonality (°C)
Min temperature (°C)
Max temperature (°C)
Water vapor pressure (kPa)
Wind speed (m/s)
Solar radiation (kJ/m²/day)
Kalahari Desert
Richtersveld
Sahara Ksar Ghilane
Sahara Douz−Matmata
Negev Desert
Kuwait Desert
North Australian Desert
South Australian Desert
Mojave Desert
Sonoran Desert
Atacama Desert
Namib Desert
Gobi China
Gobi Mongolia
Drass Valley
Mount Seuss
McMurdo Dry Valleys
Lagoon Island
Livingston Island
Hietatievat
Svalbard
Cambridge Bay
Alexandra Fiord
Kap Hoegh
Peary Land
B
pH
Gravel (%)
Sand (%)
Soil moisture (%)
Organic carbon (%)
Organic δ13C (‰)
Total δ13C (‰)
Total nitrogen (%)
Total δ15N (‰)
Ammonium (mg/kg N)
Nitrate (mg/kg N)
Sulfur (mg/kg S)
Colwell phosphorus (mg/kg P)
C
Electrical conductivity (dS/m)
ECEC (cmol+/kg)
Aluminium (%)
Boron (mg/kg)
Calcium (%)
Copper (mg/kg)
Iron (mg/kg)
Magnesium (%)
Manganese (%)
Potassium (%)
Sodium − ESP (%)
Silicon (mg/kg Si)
Zinc (mg/kg)
D
16S rRNA gene copy number
16S rRNA gene qPCR (copies/g)
Cell count
H2 oxidizer (cell/g)
CO oxidizer (cell/g)
CH4 oxidizer (cell/g)
Ammonium oxidizer (cell/g)
Nitrite oxidizer (cell/g)
Photoheterotroph (rhodopsin) (cell/g)
Photoautotroph (cell/g)
Autotroph (CBB cycle) (cell/g)
Autotroph (4−HP cycle) (cell/g)
Autotroph (rTCA cycle) (cell/g)
E
Alexandra Fiord
Atacama Desert
Cambridge Bay
Drass Valley
Gobi China
Gobi Mongolia
Hietatievat
Kalahari Desert
Kap Hoegh
Kuwait Desert
Lagoon Island
Livingston Island
McMurdo Dry Valleys
Mojave Desert
Mount Seuss
Namib Desert
Negev Desert
North Australian Desert
Peary Land
Richtersveld
Sahara Douz−Matmata
Sahara Ksar Ghilane
Sonoran Desert
South Australian Desert
Svalbard
BIO12
BIO16
BIO13
BIO14
BIO19
BIO17
BIO18
Aridity index
BIO6
BIO9
BIO11
BIO8
BIO1
WVP
BIO3
BIO10
BIO5
ET
BIO2
Elevation
Radiation
ET.Std
BIO15
Wind
BIO7
BIO4
PC1, 51.37% variation
PC2, 19.08% variation
F
Soil nitrogen
Colwell phosphorus
Nitrate
Ammonium
Soil carbon
Soil δ15N
Copper
Boron
Silicon
Organic carbon
Iron
Soil δ13C
Manganese
Moisture
Effective cation exchange capacity
Conductivity
Silt/Clay
Calcium
Sulfur
Organic δ13C
Aluminium
Sand
Gravel
Sodium
pH
Potassium
Magnesium
PC1, 26.05% variation
PC2, 16.79% variation

**Figure S2. Global soil community profiling based on small subunit RNA (SSU RNA) gene and *rplB* and zeta diversity analysis based on universal single copy ribosomal markers *rplB* and *rplP*.** (**A**) Boxplot showing the distribution of the SSU RNA gene read proportions assigned to archaea, bacteria, amoebozoa, metazoan, fungi, embryophyta, other eukaryotes, mitochondria, and chloroplast across metagenomes of different soil ecosystems by phyloFlash. (**B**) Phylum level structures of archaeal and bacterial community across different soil ecosystems by phyloFlash. Reads were taxonomically annotated by Silva database release 138. Phyla that do not attain more than 3.5% abundance in either sample were grouped to "Other phyla". (**C**) Boxplot showing relative abundance of dominant archaeal and bacterial phyla across different soil ecosystems based on *rplB*. Bacterial and archaeal taxonomy is based on the Genome Taxonomy Database. Phyla that do not attain more than 5% abundance in either sample were grouped to "Other phyla". (**D, E**) Rarefaction curves based on observed (**D**) nearest taxonomic units of the archaeal and bacterial SSU RNA gene reads (genus level) and (**E**) operational taxonomic units (genus level) of *rplB* reads in metagenomes of each site. (**F, G**) Zeta decline showing the pattern of shared richness (Jaccard normalization) of (**F**) *rplB* and (**G**) *rplP* when increasing number of sites from the same ecosystem category is considered. For each marker gene, the analysis on unrarefied dataset and datasets rarefied at 300, 400, and 500 counts at genus, family, and order taxonomic levels were shown. The enclosed boxplot shows pairwise community dissimilarity (beta diversity) between sites from the same ecosystem category using the weighted UniFrac. (**H, I**) Distance decay of zeta diversity of unrarefied and rarefied (at 400 counts) datasets of (**H**) *rplB* and (**I**) *rplP* at genus level. Pairwise (zeta order 2) and multisite (zeta orders 3 to 6) comparisons of each ecosystem were shown, with regression lines showing the generalized linear regression of shared richness (Jaccard normalization) against mean distance (km) between sites and shadings representing the 95% confidence intervals of the regression.

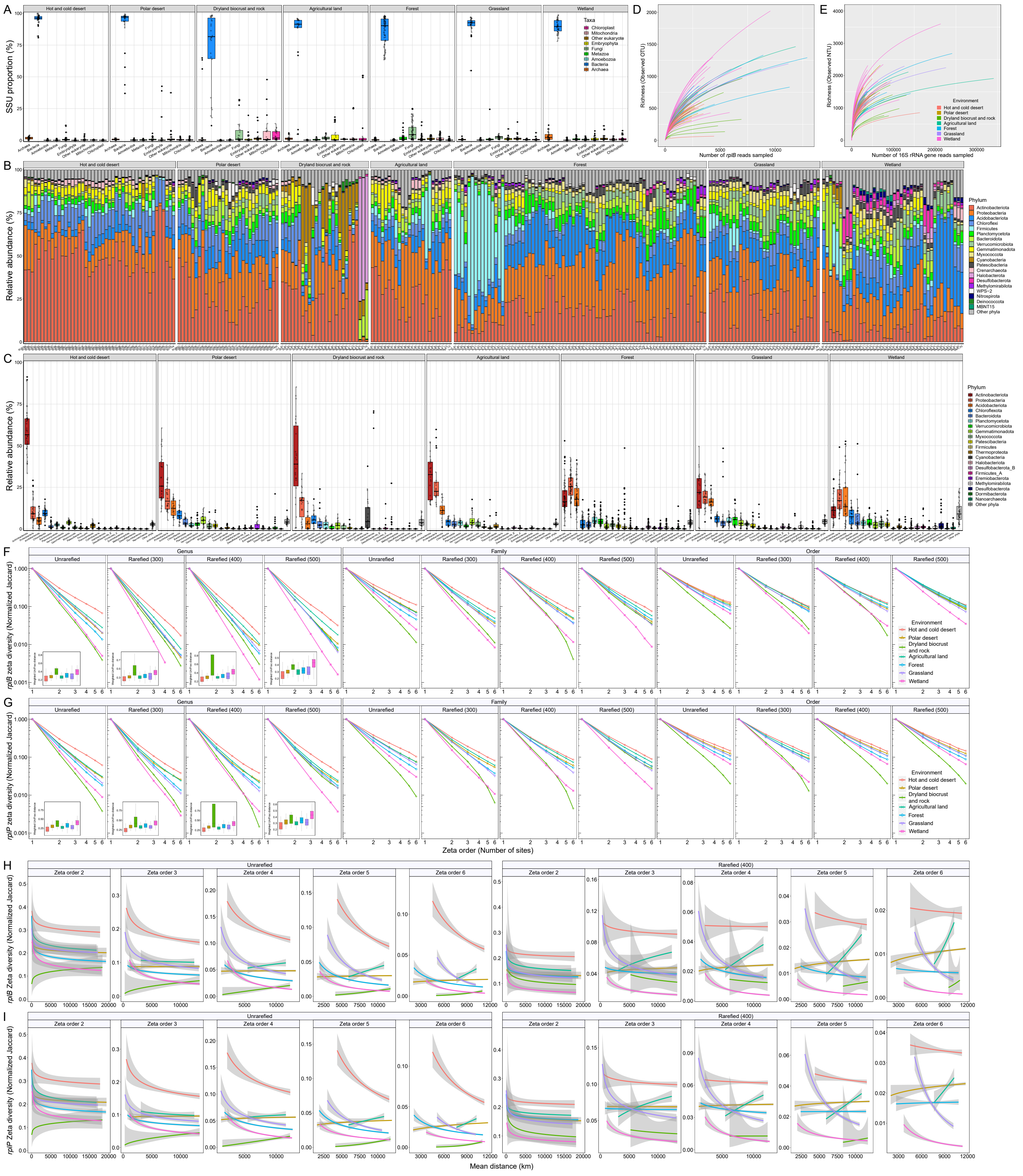
A
Hot and cold desert
Polar desert
Dryland biocrust and rock
Agricultural land
Forest
Grassland
Wetland
SSU proportion (%)
Taxa
Chloroplast
Mitochondria
Other eukaryote
Embryophyta
Fungi
Metazoa
Amoebozoa
Bacteria
Archaea
D
Richness (Observed OTU)
Number of rplB reads sampled
E
Richness (Observed NTU)
Number of 16S rRNA gene reads sampled
Environment
Hot and cold desert
Polar desert
Dryland biocrust and rock
Agricultural land
Forest
Grassland
Wetland
B
Relative abundance (%)
Phylum
Actinobacteriota
Proteobacteria
Acidobacteriota
Chloroflexi
Firmicutes
Planctomycetota
Bacteroidota
Verrucomicrobiota
Gemmatimonadota
Myxococcota
Cyanobacteria
Patescibacteria
Crenarchaeota
Halobacterota
Desulfobacterota
Methylomirabilota
WPS-2
Nitrospirota
Deinococcota
MBNT15
Other phyla
C
Relative abundance (%)
Phylum
Actinobacteriota
Proteobacteria
Acidobacteriota
Chloroflexota
Bacteroidota
Planctomycetota
Verrucomicrobiota
Gemmatimonadota
Myxococcota
Patescibacteria
Firmicutes
Thermoproteota
Cyanobacteria
Halobacteriota
Desulfobacterota_B
Firmicutes_A
Eremiobacterota
Methylomirabilota
Desulfobacterota
Dormibacterota
Nanoarchaeota
Other phyla
F
Genus
Family
Order
Unrarefied
Rarefied (300)
Rarefied (400)
Rarefied (500)
rplB zeta diversity (Normalized Jaccard)
Weighted UniFrac distance
Environment
Hot and cold desert
Polar desert
Dryland biocrust and rock
Agricultural land
Forest
Grassland
Wetland
G
rplP zeta diversity (Normalized Jaccard)
Zeta order (Number of sites)
H
Unrarefied
Rarefied (400)
Zeta order 2
Zeta order 3
Zeta order 4
Zeta order 5
Zeta order 6
rplB Zeta diversity (Normalized Jaccard)
I
rplP Zeta diversity (Normalized Jaccard)
Mean distance (km)

**Figure S3. Most relatively less abundant KEGG Orthology (KO) in desert microbiome**. The top 45 most relatively less abundant KO (with an average gene copy number over 0.25 in non-desert soil microbiome) in (**A**) hot and cold desert microbiome and (**B**) polar desert microbiome compared to global non-desert soil microbiome. The lollipop plot shows the lower quartile, median, and upper quartile of the average gene copy number per genome of each KO in short read desert metagenomes. Asterisks denote the average gene copy number of the KO is less abundant in both nonpolar and polar desert communities compared to global non-desert communities. The donut charts show the fold difference of each KO (left: short read data; right: MAGs) in nonpolar deserts and polar lands against other non-desert ecosystems. (**C**) Density plots show the distribution of the average gene copy number per genome of KO of the top 12 KEGG BRITE categories that contribute to the largest reduction in KO gene copy numbers in desert microbiome versus non-desert ecosystems.

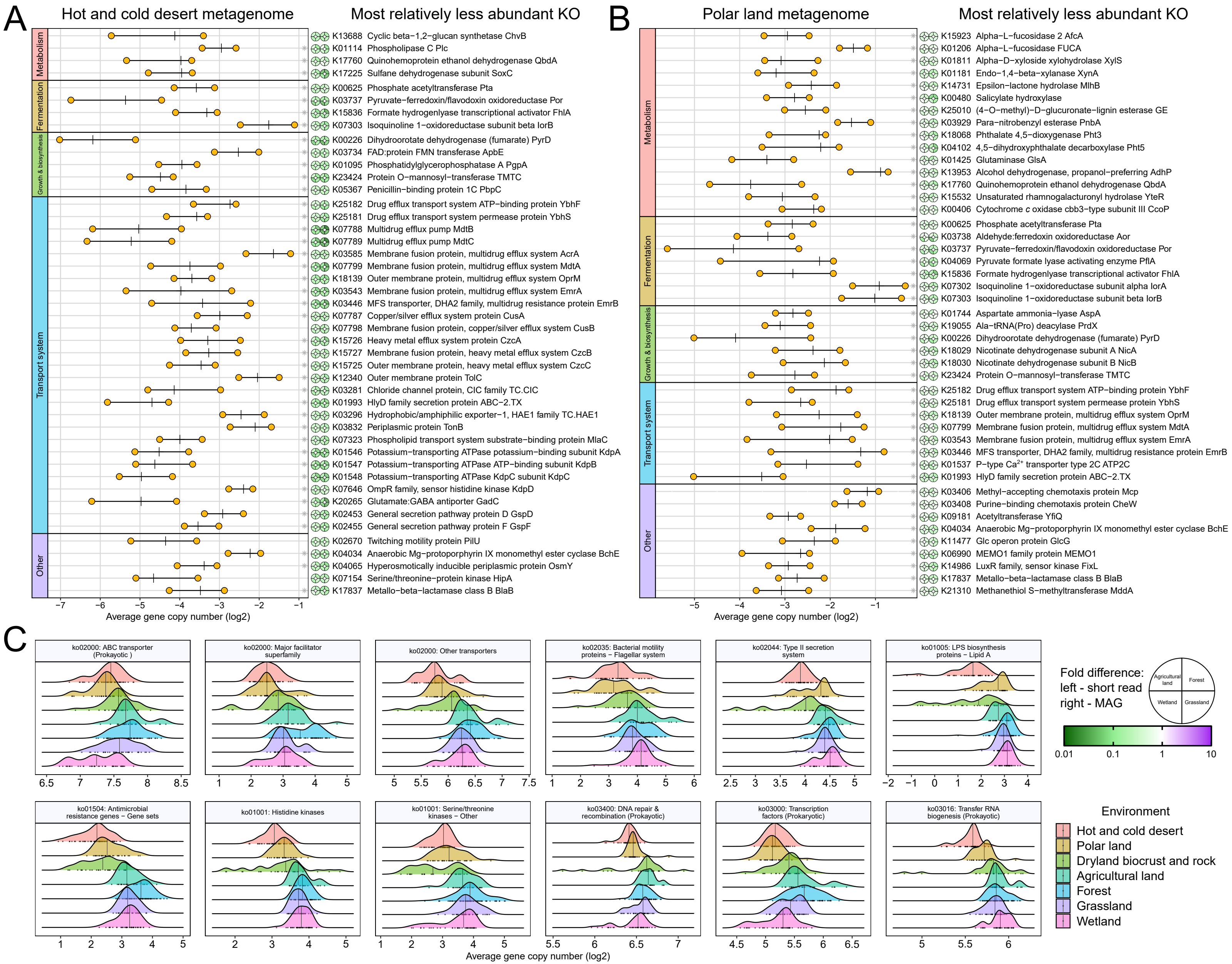
A
Hot and cold desert metagenome
Most relatively less abundant KO
Metabolism
Fermentation
Growth & biosynthesis
Transport system
Other
K13688 Cyclic beta−1,2−glucan synthetase ChvB
K01114 Phospholipase C Plc
K17760 Quinohemoprotein ethanol dehydrogenase QbdA
K17225 Sulfane dehydrogenase subunit SoxC
K00625 Phosphate acetyltransferase Pta
K03737 Pyruvate−ferredoxin/flavodoxin oxidoreductase Por
K15836 Formate hydrogenlyase transcriptional activator FhlA
K07303 Isoquinoline 1−oxidoreductase subunit beta IorB
K00226 Dihydroorotate dehydrogenase (fumarate) PyrD
K03734 FAD:protein FMN transferase ApbE
K01095 Phosphatidylglycerophosphatase A PgpA
K23424 Protein O−mannosyl−transferase TMTC
K05367 Penicillin−binding protein 1C PbpC
K25182 Drug efflux transport system ATP−binding protein YbhF
K25181 Drug efflux transport system permease protein YbhS
K07788 Multidrug efflux pump MdtB
K07789 Multidrug efflux pump MdtC
K03585 Membrane fusion protein, multidrug efflux system AcrA
K07799 Membrane fusion protein, multidrug efflux system MdtA
K18139 Outer membrane protein, multidrug efflux system OprM
K03543 Membrane fusion protein, multidrug efflux system EmrA
K03446 MFS transporter, DHA2 family, multidrug resistance protein EmrB
K07787 Copper/silver efflux system protein CusA
K07798 Membrane fusion protein, copper/silver efflux system CusB
K15726 Heavy metal efflux system protein CzcA
K15727 Membrane fusion protein, heavy metal efflux system CzcB
K15725 Outer membrane protein, heavy metal efflux system CzcC
K12340 Outer membrane protein TolC
K03281 Chloride channel protein, CIC family TC.CIC
K01993 HlyD family secretion protein ABC−2.TX
K03296 Hydrophobic/amphiphilic exporter−1, HAE1 family TC.HAE1
K03832 Periplasmic protein TonB
K07323 Phospholipid transport system substrate−binding protein MlaC
K01546 Potassium−transporting ATPase potassium−binding subunit KdpA
K01547 Potassium−transporting ATPase ATP−binding subunit KdpB
K01548 Potassium−transporting ATPase KdpC subunit KdpC
K07646 OmpR family, sensor histidine kinase KdpD
K20265 Glutamate:GABA antiporter GadC
K02453 General secretion pathway protein D GspD
K02455 General secretion pathway protein F GspF
K02670 Twitching motility protein PilU
K04034 Anaerobic Mg−protoporphyrin IX monomethyl ester cyclase BchE
K04065 Hyperosmotically inducible periplasmic protein OsmY
K07154 Serine/threonine−protein kinase HipA
K17837 Metallo−beta−lactamase class B BlaB
−7 −6 −5 −4 −3 −2 −1
Average gene copy number (log2)
B
Polar land metagenome
Most relatively less abundant KO
K15923 Alpha−L−fucosidase 2 AfcA
K01206 Alpha−L−fucosidase FUCA
K01811 Alpha−D−xyloside xylohydrolase XylS
K01181 Endo−1,4−beta−xylanase XynA
K14731 Epsilon−lactone hydrolase MlhB
K00480 Salicylate hydroxylase
K25010 (4−O−methyl)−D−glucuronate−lignin esterase GE
K03929 Para−nitrobenzyl esterase PnbA
K18068 Phthalate 4,5−dioxygenase Pht3
K04102 4,5−dihydroxyphthalate decarboxylase Pht5
K01425 Glutaminase GlsA
K13953 Alcohol dehydrogenase, propanol−preferring AdhP
K17760 Quinohemoprotein ethanol dehydrogenase QbdA
K15532 Unsaturated rhamnogalacturonyl hydrolase YteR
K00406 Cytochrome c oxidase cbb3−type subunit III CcoP
K00625 Phosphate acetyltransferase Pta
K03738 Aldehyde:ferredoxin oxidoreductase Aor
K03737 Pyruvate−ferredoxin/flavodoxin oxidoreductase Por
K04069 Pyruvate formate lyase activating enzyme PflA
K15836 Formate hydrogenlyase transcriptional activator FhlA
K07302 Isoquinoline 1−oxidoreductase subunit alpha IorA
K07303 Isoquinoline 1−oxidoreductase subunit beta IorB
K01744 Aspartate ammonia−lyase AspA
K19055 Ala−tRNA(Pro) deacylase PrdX
K00226 Dihydroorotate dehydrogenase (fumarate) PyrD
K18029 Nicotinate dehydrogenase subunit A NicA
K18030 Nicotinate dehydrogenase subunit B NicB
K23424 Protein O−mannosyl−transferase TMTC
K25182 Drug efflux transport system ATP−binding protein YbhF
K25181 Drug efflux transport system permease protein YbhS
K18139 Outer membrane protein, multidrug efflux system OprM
K07799 Membrane fusion protein, multidrug efflux system MdtA
K03543 Membrane fusion protein, multidrug efflux system EmrA
K03446 MFS transporter, DHA2 family, multidrug resistance protein EmrB
K01537 P−type Ca2+ transporter type 2C ATP2C
K01993 HlyD family secretion protein ABC−2.TX
K03406 Methyl−accepting chemotaxis protein Mcp
K03408 Purine−binding chemotaxis protein CheW
K09181 Acetyltransferase YfiQ
K04034 Anaerobic Mg−protoporphyrin IX monomethyl ester cyclase BchE
K11477 Glc operon protein GlcG
K06990 MEMO1 family protein MEMO1
K14986 LuxR family, sensor kinase FixL
K17837 Metallo−beta−lactamase class B BlaB
K21310 Methanethiol S−methyltransferase MddA
−5 −4 −3 −2 −1
Average gene copy number (log2)
C
ko02000: ABC transporter (Prokayotic )
ko02000: Major facilitator superfamily
ko02000: Other transporters
ko02035: Bacterial motility proteins − Flagellar system
ko02044: Type II secretion system
ko01005: LPS biosynthesis proteins − Lipid A
ko01504: Antimicrobial resistance genes − Gene sets
ko01001: Histidine kinases
ko01001: Serine/threonine kinases − Other
ko03400: DNA repair & recombination (Prokayotic)
ko03000: Transcription factors (Prokayotic)
ko03016: Transfer RNA biogenesis (Prokayotic)
Average gene copy number (log2)
Fold difference:
left - short read
right - MAG
Agricultural land
Forest
Wetland
Grassland
0.01 0.1 1 10
Environment
Hot and cold desert
Polar land
Dryland biocrust and rock
Agricultural land
Forest
Grassland
Wetland

**Figure S4. KEGG functional annotation of desert and global soil metagenomes. (A)** COG functional categorization of metagenomic short read annotated to KEGG database. (**B**) Abundance of annotated reads assigned to the category "Mobilome: prophages, transposons" (count per million). (**C, D**) Enriched KEGG BRITE categories in desert microbiome. Density plots show the distribution of the average gene copy number of KO assigned to (**C**) prokaryotic defence systems and (**D**) the top 36 most relatively abundant ABC transporter systems in desert microbiome compared to non-desert soil microbiome.

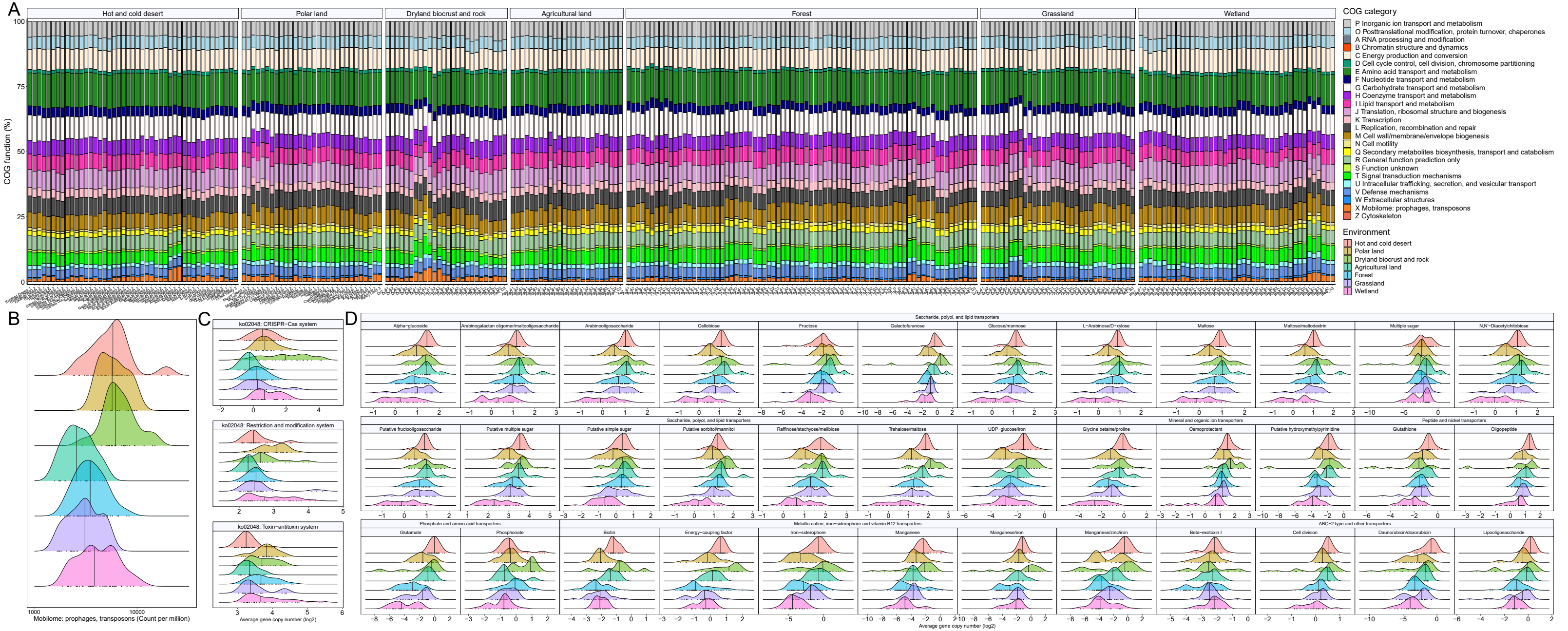
A
B
C
D
Hot and cold desert
Polar land
Dryland biocrust and rock
Agricultural land
Forest
Grassland
Wetland
COG function (%)
COG category
P Inorganic ion transport and metabolism
O Posttranslational modification, protein turnover, chaperones
A RNA processing and modification
B Chromatin structure and dynamics
C Energy production and conversion
D Cell cycle control, cell division, chromosome partitioning
E Amino acid transport and metabolism
F Nucleotide transport and metabolism
G Carbohydrate transport and metabolism
H Coenzyme transport and metabolism
I Lipid transport and metabolism
J Translation, ribosomal structure and biogenesis
K Transcription
L Replication, recombination and repair
M Cell wall/membrane/envelope biogenesis
N Cell motility
Q Secondary metabolites biosynthesis, transport and catabolism
R General function prediction only
S Function unknown
T Signal transduction mechanisms
U Intracellular trafficking, secretion, and vesicular transport
V Defense mechanisms
W Extracellular structures
X Mobilome: prophages, transposons
Z Cytoskeleton
Environment
Hot and cold desert
Polar land
Dryland biocrust and rock
Agricultural land
Forest
Grassland
Wetland
Mobilome: prophages, transposons (Count per million)
ko02048: CRISPR−Cas system
ko02048: Restriction and modification system
ko02048: Toxin−antitoxin system
Average gene copy number (log2)
Saccharide, polyol, and lipid transporters
Alpha−glucoside
Arabinogalactan oligomer/maltooligosaccharide
Arabinooligosaccharide
Cellobiose
Fructose
Galactofuranose
Glucose/mannose
L−Arabinose/D−xylose
Maltose
Maltose/maltodextrin
Multiple sugar
N,N'−Diacetylchitobiose
Putative fructooligosaccharide
Putative multiple sugar
Putative simple sugar
Putative sorbitol/mannitol
Raffinose/stachyose/melibiose
Trehalose/maltose
UDP−glucose/iron
Mineral and organic ion transporters
Glycine betaine/proline
Osmoprotectant
Putative hydroxymethylpyrimidine
Peptide and nickel transporters
Glutathione
Oligopeptide
Phosphate and amino acid transporters
Glutamate
Phosphonate
Metallic cation, iron−siderophore and vitamin B12 transporters
Biotin
Energy−coupling factor
Iron−siderophore
Manganese
Manganese/iron
Manganese/zinc/iron
ABC−2 type and other transporters
Beta−exotoxin I
Cell division
Daunorubicin/doxorubicin
Lipooligosaccharide
Average gene copy number (log2)

**Figure S5. Fraction of genome encoding carbohydrate active enzymes (CAZy) and peptidases in MAGs across different soil ecosystems**. Bar chart showing mean genome proportion of open reading frames predicted as (**A**) glycoside hydrolases, polysaccharide lyases, carbohydrate esterases, auxiliary activity proteins, and carbohydrate-binding modules, (**B**) aspartic peptidases, cysteine peptidases, metallopeptidases, serine peptidases, and periplasmic / extracellular located peptidases in MAGs from hot and cold deserts (n = 333), polar deserts (n = 578), dryland biocrust and rock (n = 528), agricultural land (n = 119), forest (n = 499), grassland (n = 386), and wetland (n = 694) (mean ± SEM). Genome proportion of the gene family was calculated by dividing the total nucleotide length of the gene against genome size of the MAG. Peptidase subcellular localization was predicted by PSORTb.

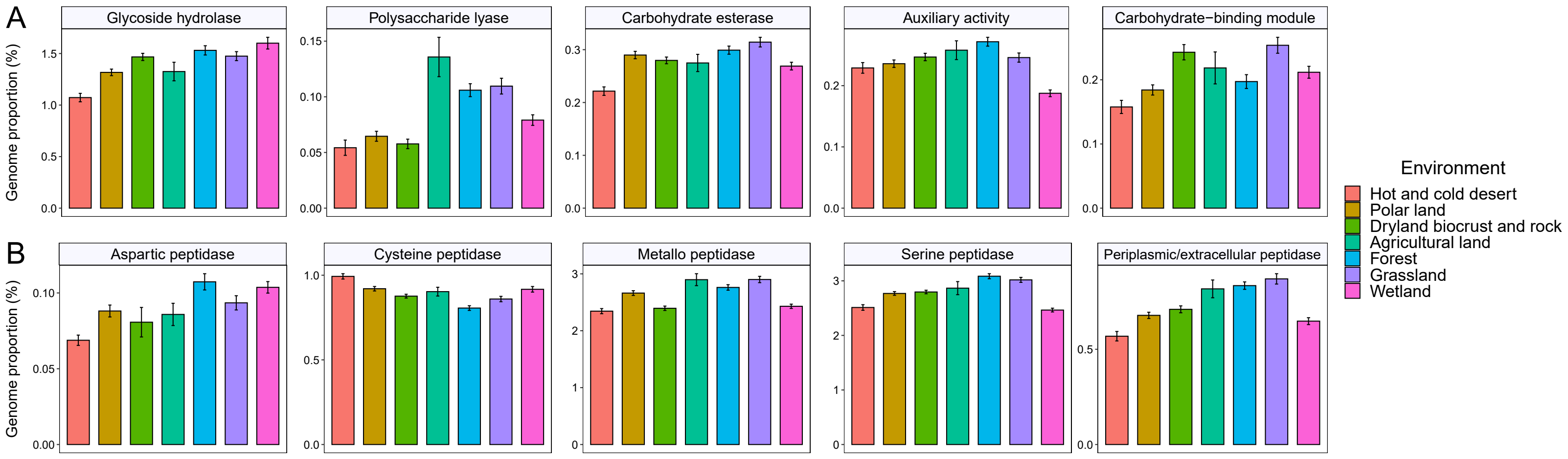
A
Glycoside hydrolase
Polysaccharide lyase
Carbohydrate esterase
Auxiliary activity
Carbohydrate−binding module
Genome proportion (%)
B
Aspartic peptidase
Cysteine peptidase
Metallo peptidase
Serine peptidase
Periplasmic/extracellular peptidase
Genome proportion (%)
Environment
Hot and cold desert
Polar land
Dryland biocrust and rock
Agricultural land
Forest
Grassland
Wetland

**Figure S6. Presence of metabolic marker genes and genome attributes of metagenome-assembled genomes from global deserts (n = 911) and non-desert soil ecosystems (n = 2226)**. (**A**) Heatmap showing the presence (%) of metabolic marker genes mediating carbon fixation, trace gas cycling, nitrogen metabolism, sulfur metabolism, arsenite, iron and formate oxidation, fumarate, iron, and selenate reduction, reductive dehalogenation, photophosphorylation, energy reserve synthesis, and electron transport chain in 911 desert MAGs. The lower panel shows the number of MAGs in each order. (**B**) Bar chart showing the presence (%) of each metabolic marker genes in MAGs from hot and cold deserts (n = 333), polar deserts (n = 578), dryland biocrust and rock (n = 528), agricultural land (n = 119), forest (n = 499), grassland (n = 386), and wetland (n = 694). (**C**) Boxplots showing the distribution of GC content, estimated completeness and contamination levels of MAGs from each ecosystem.

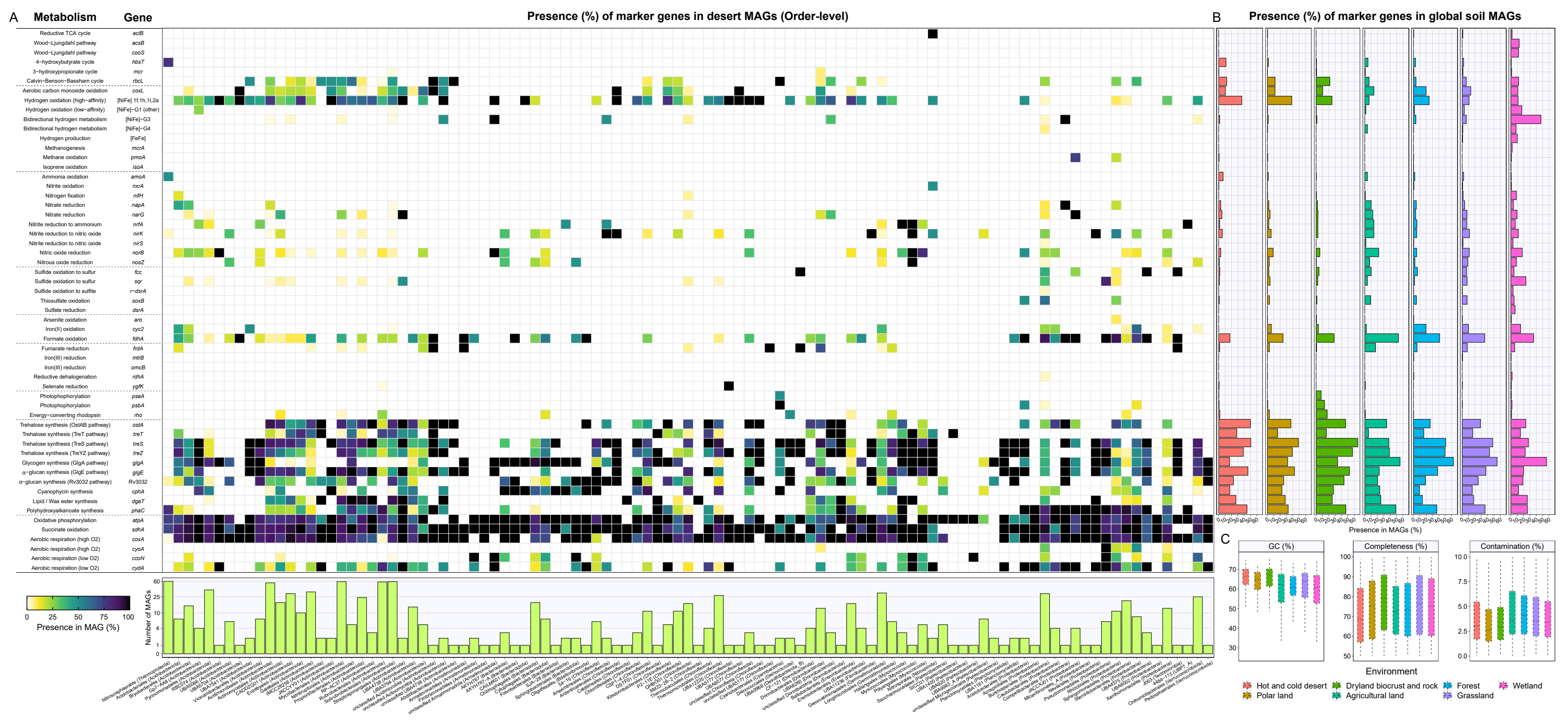

A
Metabolism
Gene
Presence (%) of marker genes in desert MAGs (Order-level)
Reductive TCA cycle
aclB
Wood−Ljungdahl pathway
acsB
Wood−Ljungdahl pathway
cooS
4−hydroxybutyrate cycle
hbsT
3−hydroxypropionate cycle
mcr
Calvin−Benson−Bassham cycle
rbcL
Aerobic carbon monoxide oxidation
coxL
Hydrogen oxidation (high−affinity)
[NiFe] 1f,1h,1l,2a
Hydrogen oxidation (low−affinity)
[NiFe]−G1 (other)
Bidirectional hydrogen metabolism
[NiFe]−G3
Bidirectional hydrogen metabolism
[NiFe]−G4
Hydrogen production
[FeFe]
Methanogenesis
mcrA
Methane oxidation
pmoA
Isoprene oxidation
isoA
Ammonia oxidation
amoA
Nitrite oxidation
nxrA
Nitrogen fixation
nifH
Nitrate reduction
napA
Nitrate reduction
narG
Nitrite reduction to ammonium
nrfA
Nitrite reduction to nitric oxide
nirK
Nitrite reduction to nitric oxide
nirS
Nitric oxide reduction
norB
Nitrous oxide reduction
nosZ
Sulfide oxidation to sulfur
fcc
Sulfide oxidation to sulfur
sqr
Sulfide oxidation to sulfite
r−dsrA
Thiosulfate oxidation
soxB
Sulfate reduction
dsrA
Arsenite oxidation
aro
Iron(II) oxidation
cyc2
Formate oxidation
fdhA
Fumarate reduction
frdA
Iron(III) reduction
mtrB
Iron(III) reduction
omcB
Reductive dehalogenation
rdhA
Selenate reduction
ygfK
Photophosphorylation
psaA
Photophosphorylation
psbA
Energy−converting rhodopsin
rho
Trehalose synthesis (OstAB pathway)
ostA
Trehalose synthesis (TreT pathway)
treT
Trehalose synthesis (TreS pathway)
treS
Trehalose synthesis (TreYZ pathway)
treZ
Glycogen synthesis (GlgA pathway)
glgA
α−glucan synthesis (GlgE pathway)
glgE
α−glucan synthesis (Rv3032 pathway)
Rv3032
Cyanophycin synthesis
cphA
Lipid / Wax ester synthesis
dgaT
Polyhydroxyalkanoate synthesis
phaC
Oxidative phosphorylation
atpA
Succinate oxidation
sdhA
Aerobic respiration (high O2)
coxA
Aerobic respiration (high O2)
cyoA
Aerobic respiration (low O2)
ccoN
Aerobic respiration (low O2)
cydA
0
25
50
75
100
Presence in MAG (%)
Number of MAGs
B
Presence (%) of marker genes in global soil MAGs
Presence in MAGs (%)
C
GC (%)
Completeness (%)
Contamination (%)
Environment
Hot and cold desert
Polar land
Dryland biocrust and rock
Agricultural land
Forest
Grassland
Wetland

**Figure S7. Correlation of gene abundance with climatological conditions across global soil and desert metagenomes (n = 275).** (**A-D**) Scatter plots showing the distribution of Spearman correlations of each (**A**) KO with aridity index, (**B**) KO with annual mean temperature, (**C**) CAZy with aridity index, and (**D**) CAZy with annual mean temperature. Only KO and CAZy (glycoside hydrolase, polysaccharide lyase, carbohydrate esterase, auxiliary activities) with average gene copies/genome over 0.05 and 0.01, respectively, determined by gene-centric short read analysis were included. Colored points in (**A-B**) indicate the most relatively abundant KO (red) and the most relative less abundant KO (green) presented in **Fig. 2C-D** and **Fig. S3A-B**, while colored in (**C-D**) indicate different CAZy families. Dashed lines denote the thresholds of the Spearman correlation to be considered as statistically significant (adjusted $p < 0.05$; Benjamini-Hochberg correction). Note that y-axis is log2-transformed. (**E-F**) Density plots showing the distribution of Spearman correlations of each (**A**) KO and (**B**) CAZy with the eight representative climatological parameters. The result shows that abundance of most genes are highly correlated with aridity.

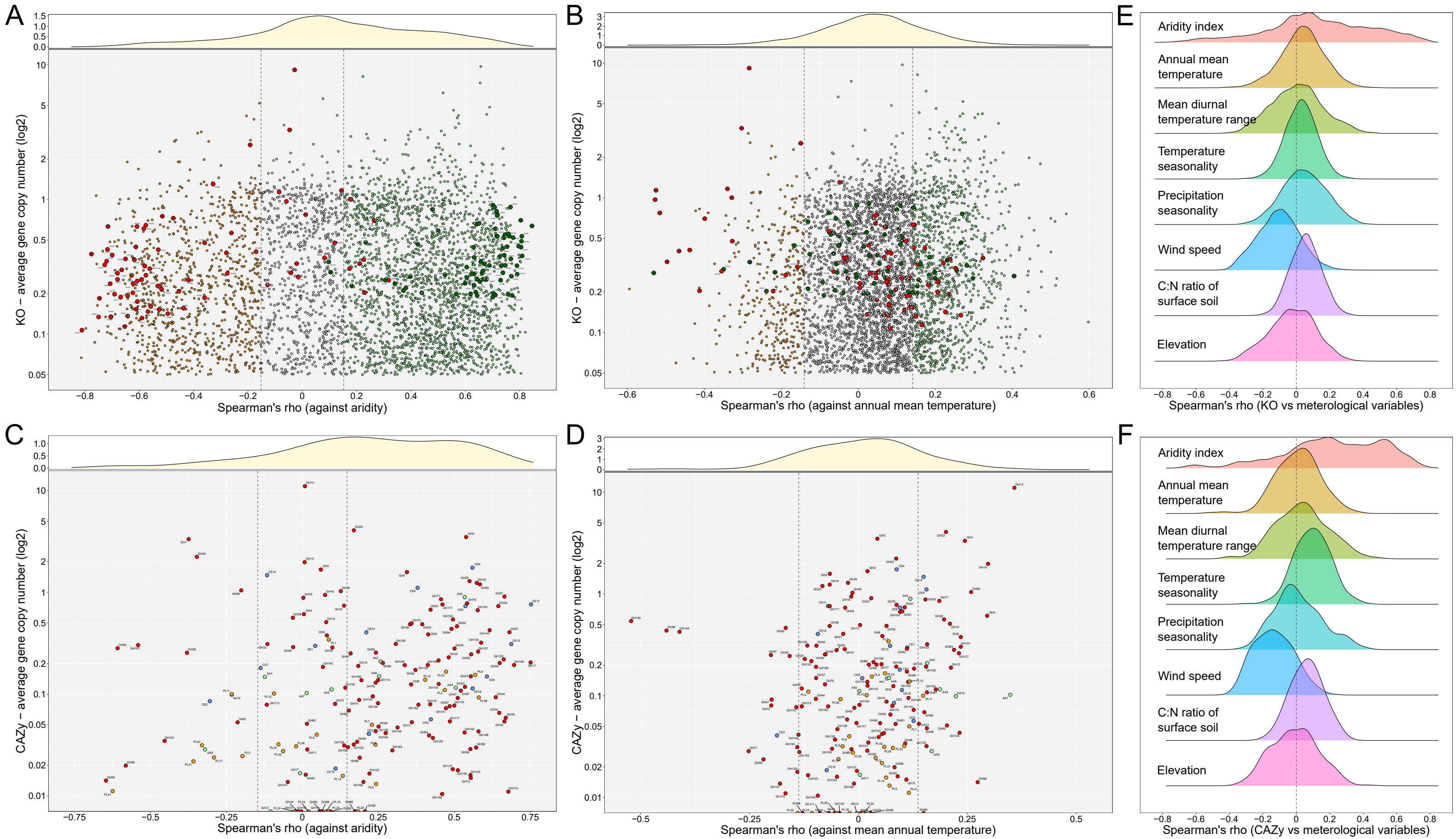
A
B
C
D
E
F
KO – average gene copy number (log2)
Spearman's rho (against aridity)
Spearman's rho (against annual mean temperature)
CAZy – average gene copy number (log2)
Spearman's rho (against aridity)
Spearman's rho (against mean annual temperature)
Aridity index
Annual mean temperature
Mean diurnal temperature range
Temperature seasonality
Precipitation seasonality
Wind speed
C:N ratio of surface soil
Elevation
Spearman's rho (KO vs meterological variables)
Spearman's rho (CAZy vs meterological variables)

**Figure S8. Gas chromatography measurements of hydrogen, carbon monoxide, and methane oxidation in desert soils**. Biological triplicate microcosms (opaque line) and heat-killed controls (translucent line) for each sampling site are shown. Hot and cold desert samples were incubated at 20°C while polar land and Drass Valley samples were incubated at 10°C. Colored lines indicate the headspace mixing ratios of $H_2$ (black), $CH_4$ (orange), and CO (blue). Solid lines and dashed lines represent trace gas measurements of soils at wetted and native hydration conditions, respectively. Under wetting, consumption to below atmospheric mixing ratios was observed for $H_2$ (all sites), $CH_4$ (Negev Desert, Mojave Desert, Sahara Douz-Matmata, South Australian Desert, Cambridge Bay, Kap Høegh, Livingston Island), and CO (all sites except Atacama Desert and Namib Desert) during the timecourse. Time is shown on a logarithmic scale.

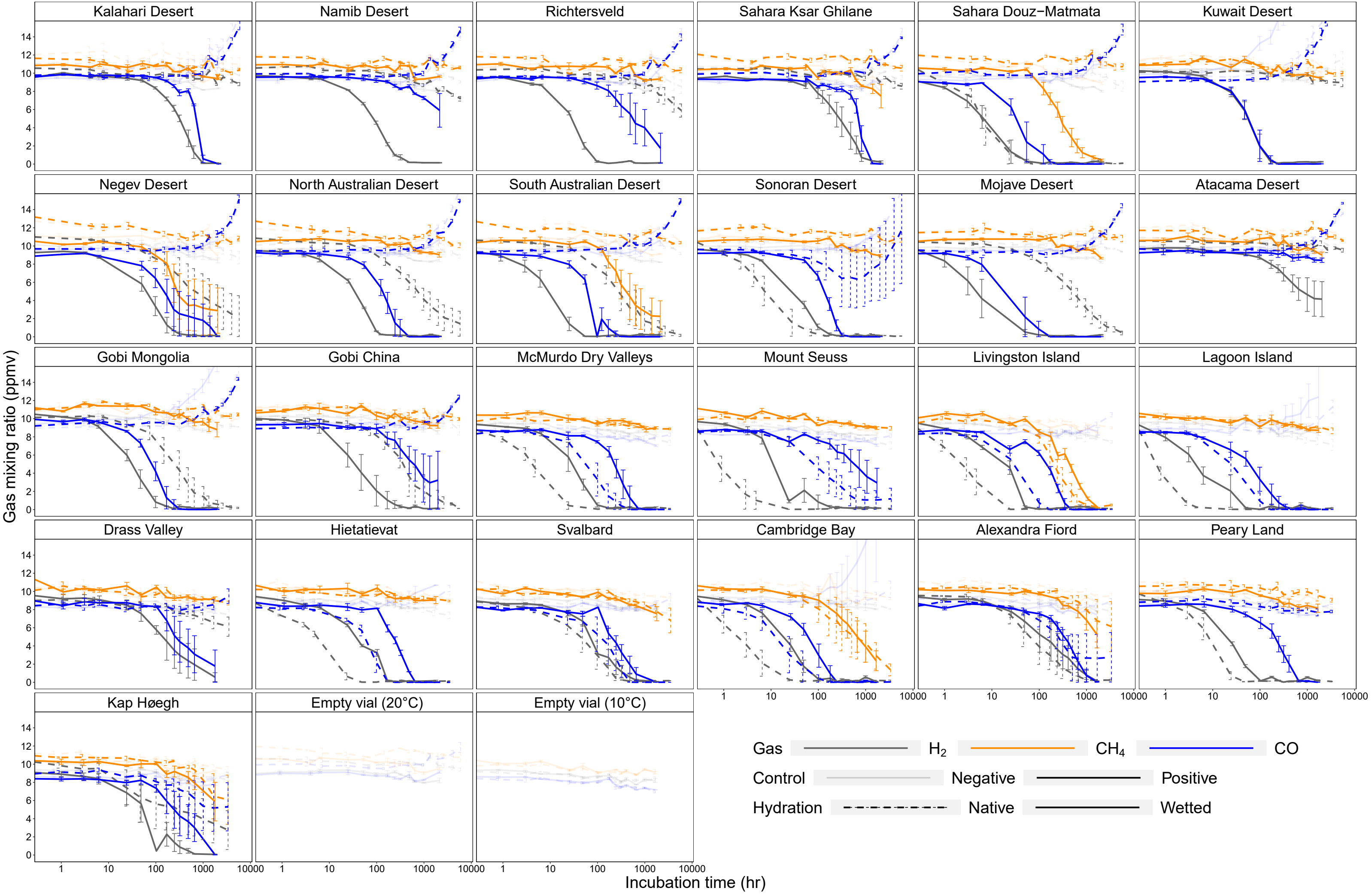
Kalahari Desert
Namib Desert
Richtersveld
Sahara Ksar Ghilane
Sahara Douz−Matmata
Kuwait Desert
Negev Desert
North Australian Desert
South Australian Desert
Sonoran Desert
Mojave Desert
Atacama Desert
Gobi Mongolia
Gobi China
McMurdo Dry Valleys
Mount Seuss
Livingston Island
Lagoon Island
Drass Valley
Hietatievat
Svalbard
Cambridge Bay
Alexandra Fiord
Peary Land
Kap Høegh
Empty vial (20°C)
Empty vial (10°C)
Gas mixing ratio (ppmv)
Incubation time (hr)
Gas
$H_2$
$CH_4$
CO
Control
Negative
Positive
Hydration
Native
Wetted

**Figure S9. $^{14}CO_2$ fixation activities of desert soils**. Rates of $^{14}CO_2$ fixation for pooled hydrated soils from each site in technical triplicate under dark ambient conditions, hydrogenotrophic conditions (100 ppmv of headspace $H_2$), and photosynthetic conditions (40 μmol photons $m^{-2}$ $s^{-1}$) (20°C for hot and cold desert samples; 10°C for polar desert and Drass Valley).

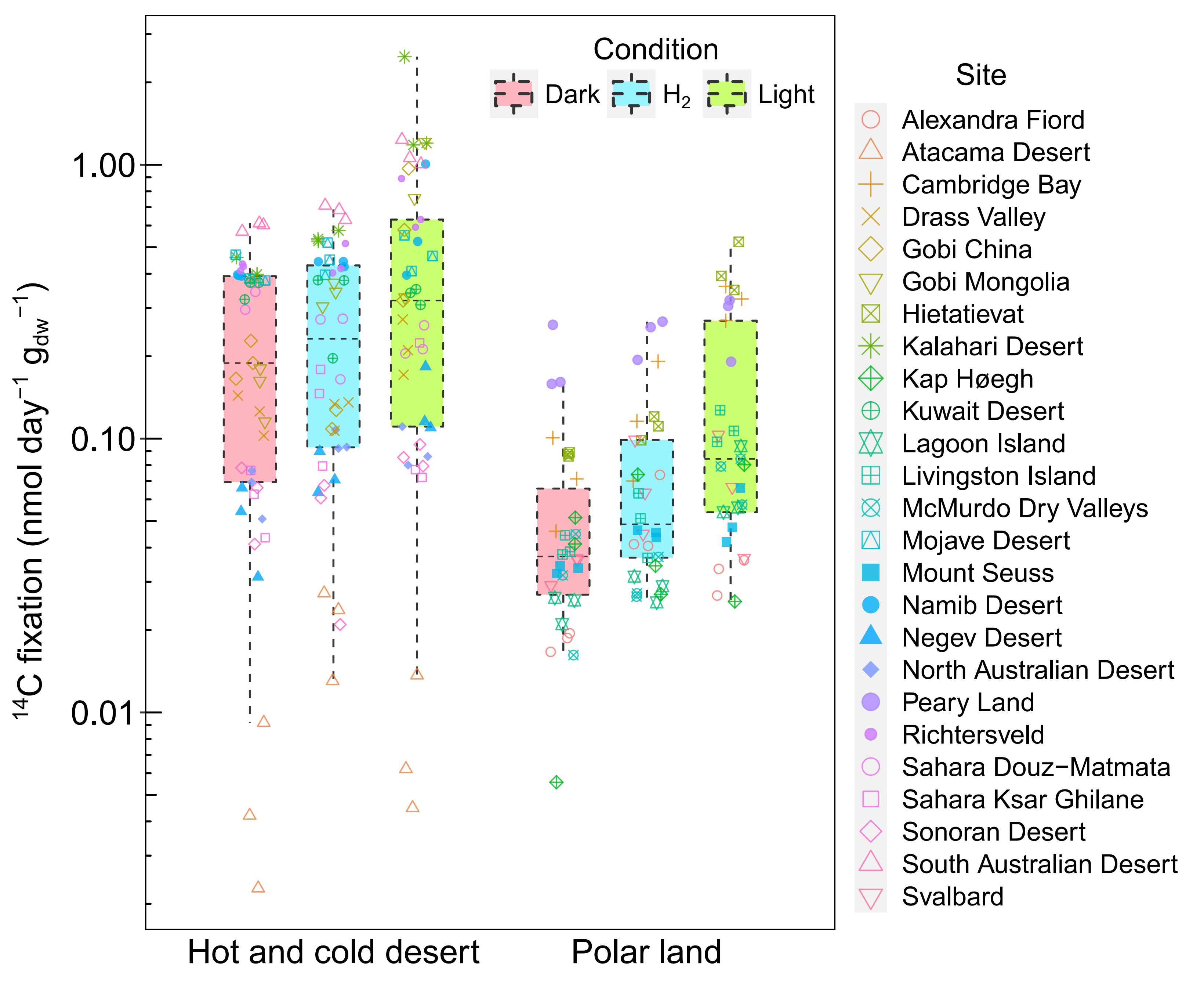

Condition
Dark
$H_2$
Light
Site
Alexandra Fiord
Atacama Desert
Cambridge Bay
Drass Valley
Gobi China
Gobi Mongolia
Hietatievat
Kalahari Desert
Kap Høegh
Kuwait Desert
Lagoon Island
Livingston Island
McMurdo Dry Valleys
Mojave Desert
Mount Seuss
Namib Desert
Negev Desert
North Australian Desert
Peary Land
Richtersveld
Sahara Douz−Matmata
Sahara Ksar Ghilane
Sonoran Desert
South Australian Desert
Svalbard
$^{14}C$ fixation (nmol $day^{-1}$ $g_{dw}^{-1}$)
1.00
0.10
0.01
Hot and cold desert
Polar land

### Supplementary Tables

**Table S1**. Sample metadata, site climate, physicochemical parameters, and 16S rRNA gene qPCR data of the 25 hot, cold, and polar desert sites.

**Table S2**. Sequencing and assembly details of the 75 desert shotgun metagenomes.

**Table S3**. Sample metadata, site climate, and metagenome information of the 200 global topsoil samples used for comparative analysis.

**Table S4**. Community composition, zeta diversity, and differential abundance analysis of soil bacterial and archaeal community using the metagenome universal single copy marker *rplB*.

**Table S5**. Community composition of soil community using the small subunit ribosomal RNA metagenomic reads.

**Table S6**. Summary of quality statistics, coverage, taxonomy, derived metabolic gene protein sequences, and genetic capabilities of metagenome-assembled genomes from deserts and global soils.

**Table S7**. Summary of KEGG annotation of short read metagenomes and metagenome-assembled genomes and significance test results.

**Table S8**. Summary of gene-centric short read metabolic analysis.

**Table S9**. Summary of random forest and Spearman correlation analyses.

**Table S10**. Measurement data for soil respiration, atmospheric $H_2$, $CH_4$, and CO oxidation, and $^{14}C$ radioisotope labelling experiments.

### References

1. M. Ortiz, *et al.*, Multiple energy sources and metabolic strategies sustain microbial diversity in Antarctic desert soils. *Proceedings of the National Academy of Sciences of the United States of America* **118**, e2025322118 (2021).

2. C. Hui, M. A. McGeoch, Zeta diversity as a concept and metric that unifies incidence-based biodiversity patterns. *American Naturalist* **184**, 684–694 (2014).

3. M. A. McGeoch, *et al.*, Measuring continuous compositional change using decline and decay in zeta diversity. *Ecology* **100** (2019).

4. M. Ji, *et al.*, Atmospheric trace gases support primary production in Antarctic desert surface soil. *Nature* **552**, 400–403 (2017).

5. S. K. Bay, *et al.*, Chemosynthetic and photosynthetic bacteria contribute differentially to primary production across a steep desert aridity gradient. *The ISME Journal* **15**, 3339–3356 (2021).

6. S. K. Bay, *et al.*, Microbial aerotrophy enables continuous primary production in diverse cave ecosystems. *bioRxiv* (2024). https://doi.org/10.1101/2024.05.30.596735.

7. H. Koch, *et al.*, Growth of nitrite-oxidizing bacteria by aerobic hydrogen oxidation. *Science* **345**, 1052–1054 (2014).

8. P. M. Leung, *et al.*, A nitrite-oxidising bacterium constitutively consumes atmospheric hydrogen. *The ISME Journal* **16**, 2213–2219 (2022).

9. G. M. King, C. F. Weber, Distribution, diversity and ecology of aerobic CO-oxidizing bacteria. *Nature Reviews Microbiology* **5**, 107–118 (2007).

10. P. R. F. Cordero, *et al.*, Atmospheric carbon monoxide oxidation is a widespread mechanism supporting microbial survival. *The ISME Journal* **13**, 2868–2881 (2019).

11. C. R. Edwards, *et al.*, Draft genome sequence of uncultured upland soil cluster Gammaproteobacteria gives molecular insights into high-affinity methanotrophy. *Genome Announcements* **5**, e00047-17 (2017).

12. S. E. Fick, R. J. Hijmans, WorldClim 2: new 1-km spatial resolution climate surfaces for global land areas. *International Journal of Climatology* **37**, 4302–4315 (2017).

13. R. J. Zomer, J. Xu, A. Trabucco, Version 3 of the global aridity index and potential evapotranspiration database. *Scientific Data* **9**, 409 (2022).
